# An extended amygdala-midbrain circuit controlling cocaine withdrawal-induced anxiety and reinstatement

**DOI:** 10.1101/2021.11.05.467532

**Authors:** Guilian Tian, May Hui, Desiree Macchia, Pieter Derdeyn, Alexandra Rogers, Elizabeth Hubbard, Chengfeng Liu, Katrina Bartas, Sean Carroll, Kevin T. Beier

## Abstract

While midbrain dopamine (DA) neuronal circuits are central to motivated behaviors, much remains unknown about our knowledge of how these circuits are modified over time by experience to facilitate selective aspects of experience-dependent plasticity. Most studies of the DA system in drug addiction focus on the role of the mesolimbic DA pathway from the ventral tegmental area (VTA) to the nucleus accumbens (NAc) in facilitating drug-associated reward. In contrast, less is known about how midbrain DA cells and associated circuits contribute to negative affective states including anxiety that emerge during protracted withdrawal from drug administration. Here, we demonstrate the selective role of a midbrain DA projection to the amygdala (VTA^DA^→Amygdala) for anxiety that develops during protracted withdrawal from cocaine administration but does not participate in cocaine reward or sensitization. Our rabies virus-mediated circuit mapping approach revealed a persistent elevation in spontaneous and task-related activity of GABAergic cells from the bed nucleus of the stria terminals (BNST) and downstream VTA^DA^→Amygdala cells that could be detected even after a single cocaine exposure. Activity in BNST^GABA^ cells was related to cocaine-induced anxiety but not reward or sensitization, and silencing the projection from these cells to the midbrain was sufficient to prevent the development of anxiety during protracted withdrawal following cocaine administration. We observed that VTA^DA^→Amygdala cells, but not other midbrain DA cells, were strongly activated after a challenge exposure to cocaine, and found that activity in these cells was necessary for the expression of reinstatement of cocaine place preference. Lastly, the importance of activity in VTA^DA^→Amygdala cells extends beyond cocaine, as these cells mediate the development of anxiety states triggered by morphine and a predator odor. Our results provide an exemplar for how to identify key circuit substrates that contribute to behavioral adaptations and reveal a critical role for BNST^GABA^→VTA^DA^→Amygdala pathway in anxiety states induced by drugs of abuse or natural experiences as well as cocaine-primed reinstatement of conditioned place preference.

## INTRODUCTION

Midbrain DA cells and their target structures such as the NAc and medial prefrontal cortex (mPFC) have been critically implicated in the development and maintenance of drug addiction. Addiction occurs in phases: initial drug exposures are rewarding; repeated administration leads to tolerance or sensitization to its effects; and withdrawal leads to anxiety and aversive phenotypes, which in turn contribute to reinstatement of drug taking/seeking^1^. Each of these adaptations is dependent on midbrain DA cells^2,3^, likely through the induction of DA-dependent plasticity in target brain structures^4^. Though midbrain DA cells were originally thought to be a largely homogenous cell population, recent studies have demonstrated that they are heterogeneous in their transcriptional and electrophysiological profiles, behavioral functions, projection sites, and input patterns^5–7^. Therefore, in order to understand how drug use and withdrawal contribute to long-lasting changes in animal behavior, a more nuanced picture of how selected midbrain DA cells contribute to specific aspects of behavioral adaptation is required.

DA is critical for reward and aversion-related behavior^6,8,9^, and modifications to VTA^DA^ cells enables processing of both positive and negative reinforcements to enact adaptation to a dynamic environment^10,11^. While the contribution of DA cells to reward and aversion learning is relatively well understood, our knowledge of how experience modifies the functional properties of these cells to facilitate learning, for example in the context of drug use, is incomplete. Drugs of abuse trigger long-lasting changes in the brain’s reward circuitry through release of DA from VTA cells in downstream structures as well as the VTA itself^16,17^. While the majority of addiction research has been focused on how drug-evoked DA release in the NAc contributes to reward associated with drug use^8,17–20^, DA is involved in a wide array of other biological processes. For example, studies have implicated DA in experience-induced anxiety, as infusion of DA D2-receptor antagonists into the amygdala attenuates anxiety induced by conditioned fear^21^. This effect is likely mediated by VTA^DA^ cells, as infusion of D2 receptor antagonists into the VTA mirrors effects of infusion into the amygdala^21,22^. In addition, elevations in DA neuron activity are linked to anxiety, as decreasing activity in VTA cells can reverse the elevation in anxiety induced by increasing DA cell activity^2^. These results together point to the potential requirement of activation of VTA^DA^→Amygdala cells for the development and expression of anxiety-related behaviors. However, whether activity of this particular cell population is necessary for the development of anxiety states, including those induced by withdrawal following repeated administration of drugs of abuse, has not been definitively demonstrated. Furthermore, the identity of input populations that control activation of these cells and how they are modified by experience is not clear. For example, while the brain’s stress system, including the extended amygdala is known to play a prominent role in drug-induced withdrawal, our understanding of how the extended amygdala and midbrain DA system may work together to coordinate anxiety-driven behaviors associated with drug withdrawal is incomplete. The recent advent of the monosynaptic rabies virus (RABV) mapping strategy has enabled the dissection of input contributions to behavior, and how selected input populations influence defined DA cell populations^9,12–15^. However, much remains to be understood about how input populations contribute to DA-dependent learning in a broad range of reward and aversion-related behaviors, including those elicited by drugs of abuse.

Here we demonstrate that VTA^DA^→Amygdala cells are critical for the development of cocaine withdrawal-induced anxiety. Activity of these cells during cocaine exposure cells was required for the subsequent development of anxiety during protracted withdrawal following repeated cocaine administration, but had no effect on cocaine-induced place preference or sensitization. Furthermore, silencing activity in DA cells projecting to the NAc or mPFC, or substantia nigra DA neurons (SNcDA) projecting to the dorsolateral striatum (DLS) during cocaine administration had no effect on the development of anxiety. We then use RABV-mediated circuit mapping, calcium imaging, and chemogenetic inhibition in defined cell types and projections to demonstrate that BNST^GABA^ cells control the development of anxiety states through their projections to the midbrain via VTA^DA^→Amygdala cells. The importance of the BNST^GABA^→VTA^DA^→Amygdala pathway extends beyond cocaine, as activity in this pathway is required for anxiety states induced by other drugs of abuse including the model opioid morphine as well as natural experiences such as predator odors. Our study highlights how our integrated activity mapping and functional circuit interrogation can illuminate fundamental principles of how experiences cause enduring changes in key brain circuits that facilitate development of basic behavioral adaptations.

## RESULTS

### Activity in VTA^DA^→Amygdala cells is selectively required for cocaine-induced anxiety

We first wanted to identify which midbrain DA cells were responsible for the development of anxiety that occurs during protracted withdrawal following repeated cocaine administration. We therefore silenced activity of unique midbrain DA cells delineated by projection to one of five projection sites: medial shell of the NAc (NAcMed), lateral shell of the NAc (NAcLat), dorsolateral striatum (DLS), amygdala, or mPFC. To target each cell population, *CAV-FLEx^loxP^-Flp,* a canine adenovirus that expresses the Flp recombinase in neurons that both 1) project to the site where CAV is injected and 2) express the Cre recombinase^12,23^, was injected into one of the five forebrain sites. During the same surgery, an adeno-associated virus (AAV) expressing Flp-dependent hM4Di, an inhibitory DREADD^24^, or YFP (*AAV_DJ_-hSyn-FLEx^FRT^-hM4Di* or *AAV_DJ_-hSyn-FLEx^FRT^-YFP*) was injected into the VTA or adjacent SNc (Fig. 1A). After allowing two weeks for expression of hM4Di or YFP, we began our protocol (Fig. 1B). In order to elicit anxiety during the abstinence period, we needed to perform multiple cocaine administrations prior to abstinence. During the cocaine administration period, we tested for the development of cocaine-induced place preference (CPP) and locomotor sensitization. CPP is a commonly used measure of cocaine reward, while locomotor sensitization is a common and robust behavioral adaptation in rodents that occurs in response to repeated administration of drugs such as cocaine. The mechanisms underlying CPP and locomotor sensitization are thought to contribute to drug-induced pathophysiological motivational states associated with addiction^25–28^. Two additional control groups were included whereby animals received injections of AAVs encoding YFP or hM4Di, but received saline injections instead of cocaine. Activity in each DA cell population was modulated by IP injection of 5 mg/kg clozapine-N-oxide (CNO) that was administered 30 min before cocaine injection (15 mg/kg). All groups received CNO. Mice then underwent ten days of protracted withdrawal. After the ten days of abstinence, we tested anxiety by both the elevated plus maze (EPM) and time spent in the center of the open field (OFT). Neither cocaine nor CNO was administered during anxiety tests as we aimed to measure baseline anxiety during protracted withdrawal.

**Figure 1:**
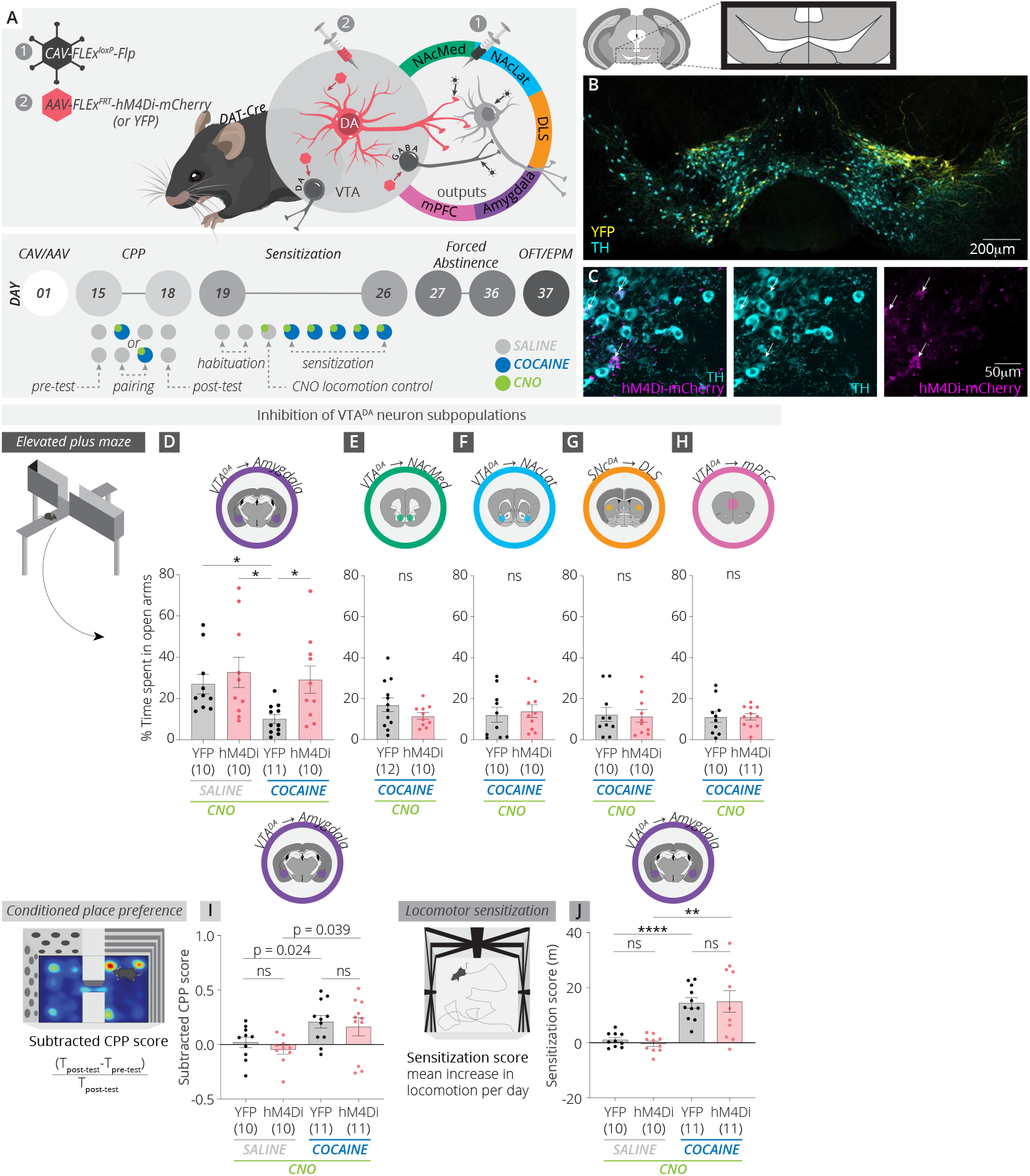
Activity in VTA^DA^→Amygdala cells is selectively required for cocaine-induced anxiety. (A) Schematic and timeline of experiments. *CAV-FLEx^loxP^-Flp* was injected into one of 5 terminal fields of DA neurons, and during the same surgery, AAVs expressing Flp-dependent hM4Di or YFP were injected into the ventral midbrain to target expression of hM4Di or YFP to output-defined DA cell subpopulations. The effects of inhibition of each subtype was tested on a variety of cocaine-induced behaviors. CPP, conditioned place preference; OFT, open field test; EPM, elevated plus maze. (B) Expression of YFP in VTA^DA^→Amygdala cells. (C) Overlap of Flp-dependent hM4Di expression with tyrosine hydroxylase, a marker of DA neurons. (D) We observed a reduction in the time spent in the open arms of the EPM during protracted withdrawal following repeated cocaine administration. This reduction was blocked by hM4Di-mediated inhibition of VTA^DA^→Amygdala cells (uncorrected p-values, YFP saline vs YFP cocaine, p = 0.0032; YFP saline vs. hM4Di saline, p = 0.53; YFP saline vs hM4Di cocaine, p = 0.79; YFP cocaine vs. hM4Di saline, p = 0.0064; YFP cocaine vs. hM4Di cocaine, p = 0.01; hM4Di saline vs. hM4Di cocaine, p = 0.73). (E-H) Inhibition of NAcMed-projecting VTA^DA^ cells (p = 0.94), NAcLat-projecting VTA^DA^ cells (p = 0.71), DLS-projecting SNcDA cells (p = 0.87), or mPFC-projecting VTA^DA^ cells (p = 0.99) had no consequences for anxiety as assessed using the EPM. (I) Inhibition of VTA^DA^→Amygdala cells during cocaine administration had no effects on cocaine CPP (uncorrected p-values, YFP saline vs YFP cocaine, p = 0.024; YFP saline vs. hM4Di saline, p = 0.31; YFP saline vs hM4Di cocaine, p = 0.16; YFP cocaine vs. hM4Di saline, p = 0.0022; YFP cocaine vs. hM4Di cocaine, p = 0.65; hM4Di saline vs. hM4Di cocaine, p = 0.39). (J) Inhibition of VTA^DA^→Amygdala cells during cocaine administration had no effects on cocaine-induced locomotor sensitization (uncorrected p-values, YFP saline vs YFP cocaine, p < 0.0001; YFP saline vs. hM4Di saline, p = 0.26; YFP saline vs hM4Di cocaine, p = 0.0035; YFP cocaine vs. hM4Di saline, p < 0.0001; YFP cocaine vs. hM4Di cocaine, p = 0.91; hM4Di saline vs. hM4Di cocaine, p = 0.0017). Error bars in this figure and throughout the manuscript = ± 1 SEM.

We found that chemogenetic inhibition of VTA^DA^→Amygdala cells during cocaine administration prevented the development of withdrawal-induced anxiety (Fig. 1D, S1). Inhibition of four other subpopulations of midbrain DA cells had no effect on anxiety (Fig. 1E-H). Interestingly, the effects were specific for the development of withdrawal anxiety, as inhibition of VTA^DA^→Amygdala cells had no effect on cocaine reward or sensitization (Fig. 1I-J). CNO administration had no effect on locomotion in animals expressing either YFP or hM4Di in VTA^DA^→Amygdala cells (Fig. S1K), a single CNO pairing for these animals in the CPP assay did not elicit CPP or CPA (Fig. S1L), and repeated administration of CNO did not trigger long-lasting anxiety phenotypes (Fig. S1M-N), demonstrating that the anxiety behavior we observed was due to withdrawal following cocaine administration, and that CNO application to hM4Di-expressing or YFP-expressing control animals did not have observable behavioral consequences in our assays. These results suggest that the rewarding effects of cocaine and psychomotor sensitization are dissociable from anxiety that develops during withdrawal, and that activity of VTA^DA^→Amygdala cells is required during cocaine administration specifically for development of anxiety that occurs during protracted withdrawal following repeated cocaine administration.

### RABV tracing reveals changes in BNST inputs to VTA^DA^→Amygdala cells

Salient experiences such as cocaine intake can trigger distinct synaptic modifications onto different projection-defined midbrain DA subtypes^29^. One limitation with previous studies is that they have mostly focused on excitatory synapses, and have lacked the resolution to identify which inputs are altered onto each DA cell subtype. In contrast, our RABV-mediated strategy enables identification of input cell populations that are modified by experience, including inputs from both GABAergic and glutamatergic cells^30^. While our previous study mapped cocaine-induced input changes onto all midbrain DA cells irrespective of projection site, here we combined our circuit screen with our output-specific viral mapping strategy, cell-type specific tracing the Relationship between Inputs and Outputs (cTRIO)^12,14,23^ to identify which input populations to VTA^DA^→Amygdala cells were modified by cocaine. Briefly, *CAV-FLEx^loxP^-Flp* was injected into the amygdala, and during the same surgery, Flp-dependent AAVs expressing the avian TVA protein tagged to mCherry (TC) and RABV glycoprotein (RABV-G) and were injected in the VTA. After waiting 13 days to enable expression of TC and RABV-G, we injected either a single dose of cocaine (15 mg/kg) or saline as a control. The following day, an EnvA-pseudotyped, G-deleted, GFP-expressing RABV (RABVΔG) was injected into the VTA. The EnvA glycoprotein binds to TVA and enables infection specifically of TC-expressing cells; therefore, the EnvA-pseudotyped RABVΔG infected only TC-expressing (and RABV-G-expressing) cells in the VTA. While the RABV was G-deleted and thus could not spread on its own, RABV-G was supplied *in trans* in the starter cells, enabling the virus to spread to connected input cells. Since the inputs did not express RABV-G, the virus could not spread further.

To test for cocaine-induced changes in inputs, we examined inputs from 22 different anatomically-defined brain regions that combined make up >90% of long-range inputs to these cells^12,30,31^. We observed RABV-labeled neurons in each examined brain site in both conditions, the majority of which had quantitatively similar proportions in saline- and cocaine-treated animals. However, we identified a significant increase in labeled inputs from the BNST onto VTA^DA^→Amygdala cells in cocaine-treated animals (saline, 1.76% of inputs; cocaine, 5.54% of inputs, p = 0.0014; Fig. 2B); this was the only brain region to reach significance, and still was significant even when correcting for multiple comparisons. These changes were specific to cocaine administration, as an aversive foot shock stimulus did not produce this increase in input labeling (Fig. S2). Interestingly, we did not observe this elevation in BNST input labeling when sampling inputs onto all VTA^DA^ cells^30^, suggesting that this increase in input labeling was specific to subpopulations of VTA^DA^ cells projecting to the amygdala.

**Figure 2:**
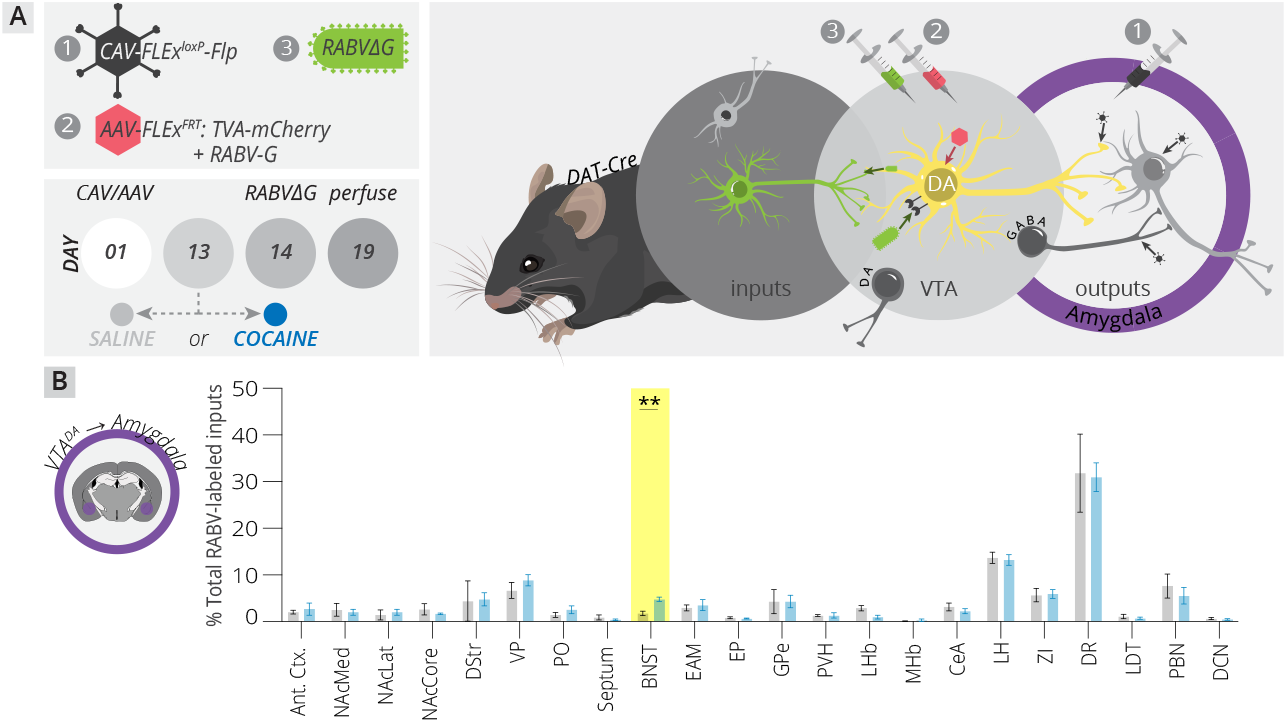
cTRIO detects an increase in BNST input labeling onto VTA^DA^→Amygdala cells in cocaine-treated mice. (A) Schematic and timeline of cTRIO experiments for VTA^DA^→Amygdala cells. *CAV-FLEx^loxP^-Flp* was injected into the amygdala, and during the same surgery, AAVs expressing Flp-dependent TC and RABV-G were injected into the VTA. Thirteen days later, animals were given a single dose of cocaine, or saline as a control. The following day, RABV-EnvA was injected into the VTA. Animals were sacrificed five days later. (B) Cocaine triggered an increase in RABV-labeled inputs from the BNST onto VTA^DA^→Amygdala cells (p = 0.0014). No other significant changes were observed.

### Activity in BNST^GABA^ cells is elevated following a single cocaine exposure

We showed previously that RABV-mediated input labeling is increased in more highly active connections, and decreased in less active ones^30^. In that study, we first linked the elevation of RABV-labeled GPe inputs to VTA^DA^ neurons with an elevation of *in vivo* and *ex vivo* activity of GPe cells. When then used chemogenetic or Kir2.1-mediated elevations or depressions in GPe cell activity to directly demonstrate that increasing or decreasing activity in these cells can elevate or depress input labeling in the targeted brain region^30^. This was true for both GPe and NAcMed inputs to VTA^DA^ cells, suggesting that it is a general property of RABV mapping. However, given that our previous observation was the first published report of this phenomenon, we wanted to test if the elevation of BNST cell labeling observed here was due to changes in BNST cellular activity, or due to another cause such as a change in the number of synapses from BNST cells in the ventral midbrain.

In order to assess whether there were changes in input synapse number or in input cell activity, we first needed to know the identity of BNST cells projecting to the midbrain. While both GABAergic and glutamatergic cells in the BNST synapse onto both VTA^DA^ and VTA^GABA^ cells^32^, the proportion of these cells that connect to VTA^DA^→Amygdala cells is not known. Therefore we used in situ hybridization in cTRIO brains mapping inputs to VTA^DA^→Amygdala cells to test if RABV-labeled inputs predominately arose from GABAergic or glutamatergic cells, and if cocaine altered the proportion of each cell type labeled. We observed that the majority of RABV-labeled BNST cells projecting to VTA^DA^→Amygdala cells were GABAergic (~75%), and that this proportion was approximately the same in both saline and cocaine-treated animals (Fig. 3A-D). Given that GABAergic cells comprised the majority of inputs from the BNST to VTA^DA^→Amygdala cells, we used the *vGAT-Flp* driver line moving forward to access these cells.

**Figure 3:**
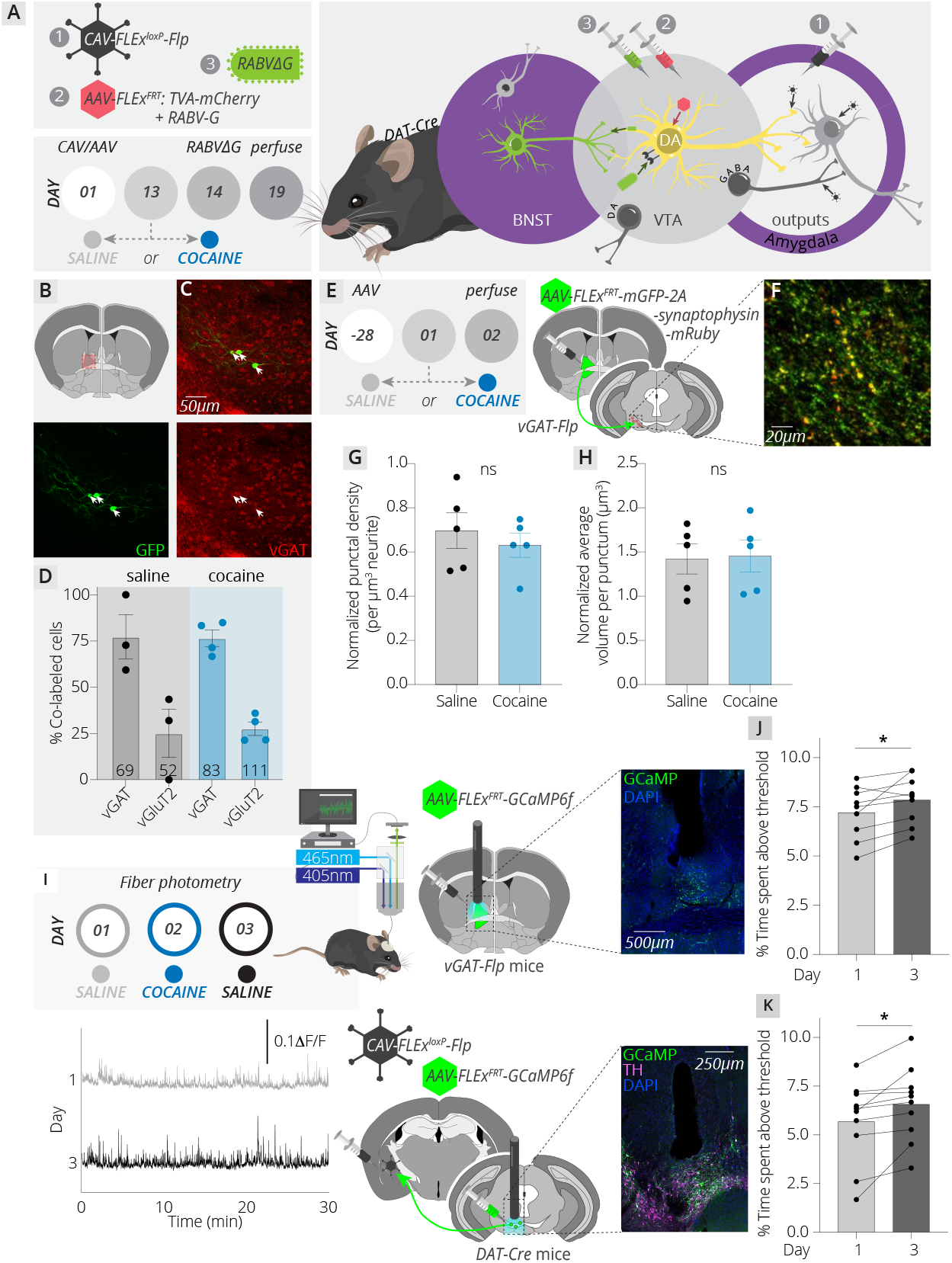
A single dose of cocaine causes long-lasting changes in the spontaneous activity of BNST^GABA^ and VTA^DA^→Amygdala cells. (A) cTRIO using the amygdala as an output site was used to label BNST cells projecting to VTA^DA^→Amygdala cells. (B) We performed fluorescence in situ hybridization (FISH) experiments, focusing on the BNST. (C) Probes to either *Slc32a1* (vGAT) or *Slc17a6* (vGluT2) were used to identify GABAergic or glutamatergic neurons, respectively. A representative image of 3 RABV-labeled neurons colocalizing with the *Slc32a1* probe is shown. (D) Quantification of overlap of RABV-labeled cells with FISH probes. The total number of cells counted for each condition is provided in the respective bar. n=3 for saline group, n = 4 for cocaine group. (E) Timeline and schematic of quantification of putative synapses from BNST^GABA^ cells in the midbrain. (F) Sample image of GFP axons and synaptophysin-mRuby puncta in the ventral midbrain. (G-H) No changes in the density (G) or volume (H) of puncta were observed. (I) Schematic and timeline of fiber photometry experiments, as well as representative traces from D1 and D3 in BNST^GABA^ cells. (J) A single dose of cocaine caused a long-lasting increase in activity of BNST^GABA^ cells (D1-D3, p = 0.04). (K) These changes corresponded to an increase in activity of VTA^DA^→Amygdala cells (D1-D3, p = 0.01).

Using this line, we first tested if cocaine altered the number of synapses from BNST^GABA^ cells in the midbrain by injecting *AAV_DJ_-hSyn-FLEx^FRT^-mGFP-2A-Synaptophysin-mRuby* into the BNST. This allowed us to quantify the number and size of putative synaptic puncta in the midbrain that arose from BNST^GABA^ cells. We observed no detectable difference in either synapse number or volume, suggesting that cocaine did not alter the numbers of inputs from these cells in the midbrain (Fig. 3E-H). To test if a single exposure to cocaine caused a long-lasting change in activity of BNST^GABA^ cells, we used fiber photometry to measure spontaneous activity of BNST^GABA^ cells one day before and one day following cocaine administration. Though these cells do project to other brain sites such as the amygdala, periaqueductal gray (PAG), and NAc, the dominant projection was to the ventral midbrain, in particular the substantia nigra pars reticulata (SNr), a brain region primarily composed of GABAergic neurons (Fig. S3). As the major projection of BNST^GABA^ cells was to the ventral midbrain, we performed fiber photometry recordings from all BNST^GABA^ cells. We found that spontaneous activity of BNST^GABA^ cells was elevated 24 hours following cocaine, reflecting the increase in RABV labeling observed following cocaine (Fig. 3I-J).

Given that the activity of BNST^GABA^ cells was elevated following cocaine, we wanted to know how this change may impact activity of downstream VTA^DA^→Amygdala cells. Though the elevation in input labeling was detected onto VTA^DA^→Amygdala cells (Fig. 2B), it was previously shown that BNST^GABA^ cells provide synaptic input to both midbrain DA and GABA cells, with a preferential input onto GABA cells^32^. This is consistent with our observation that the dominant midbrain projection of BNST^GABA^ cells is to the SNr, and not the VTA or SNc where the DA cells are located (Fig. S3). Given these observations, we expected that an elevation of BNST^GABA^ cell activity would lead to an increase in VTA^DA^→Amygdala cell activity through disinhibition via local midbrain GABA cells. Indeed, we found that activity in VTA^DA^→Amygdala cells was elevated one day following cocaine (Fig. 3K). These results are consistent with characterized connectivity of BNST^GABA^ to ventral midbrain cells, and similar to our previous results with GPe^PV^ inputs to VTA ^DA^ neurons^30^. Furthermore, they provide additional support for the notion that the net effect of activity changes in inputs on DA cell activity cannot be readily predicted from RABV mapping results alone, but rather the net effect on DA cells depends on local connectivity of microcircuits in the midbrain.

### Activity in the BNST^GABA^→midbrain projection is required for cocaine-induced anxiety

We showed that cocaine leads to elevated activity in BNST^GABA^ and VTA^DA^→Amygdala cells (Fig. 3J-K), and that activity in VTA^DA^→Amygdala cells was necessary for the development of anxiety that develops during protracted withdrawal from cocaine exposure (Fig. 1D). Given the parallel increases in activity of BNST^GABA^ and VTA^DA^→Amygdala cells, we anticipated that inhibiting the BNST^GABA^→midbrain projection during cocaine administration would also prevent the development of anxiety without impacting cocaine reward or sensitization. To test this, we injected *AAV_DJ_-hSyn-FLEx^FRT^-hM4Di* or *AAV_DJ_-hSyn-FLEx^FRT^-YFP* as a control into the BNST of *vGAT-Flp* mice, waited six weeks to enable terminal expression of hM4Di or YFP, and then began behavioral testing. In order to specifically inhibit the BNST^GABA^→midbrain projection, we injected slow-release CNO microspheres into the ventral midbrain^30,34^. These microspheres release CNO for about ~7 days; therefore, we condensed our protocol to enable administration of all six cocaine injections within this time window (Fig. 4A-B). We found that inhibition of the BNST^GABA^→midbrain projection prevented the development of anxiety during abstinence following repeated cocaine injection with no effect on cocaine reward or sensitization (Fig. 4C-E, Fig. S4), the same effect as inhibiting VTA^DA^→Amygdala cells during cocaine administration (Fig. 1D, I, J, S1). These results together demonstrate that activity in the BNST^GABA^→VTA^DA^→Amygdala pathway during cocaine administration is required for the development of anxiety that occurs following protracted withdrawal from repeated cocaine administration.

**Figure 4:**
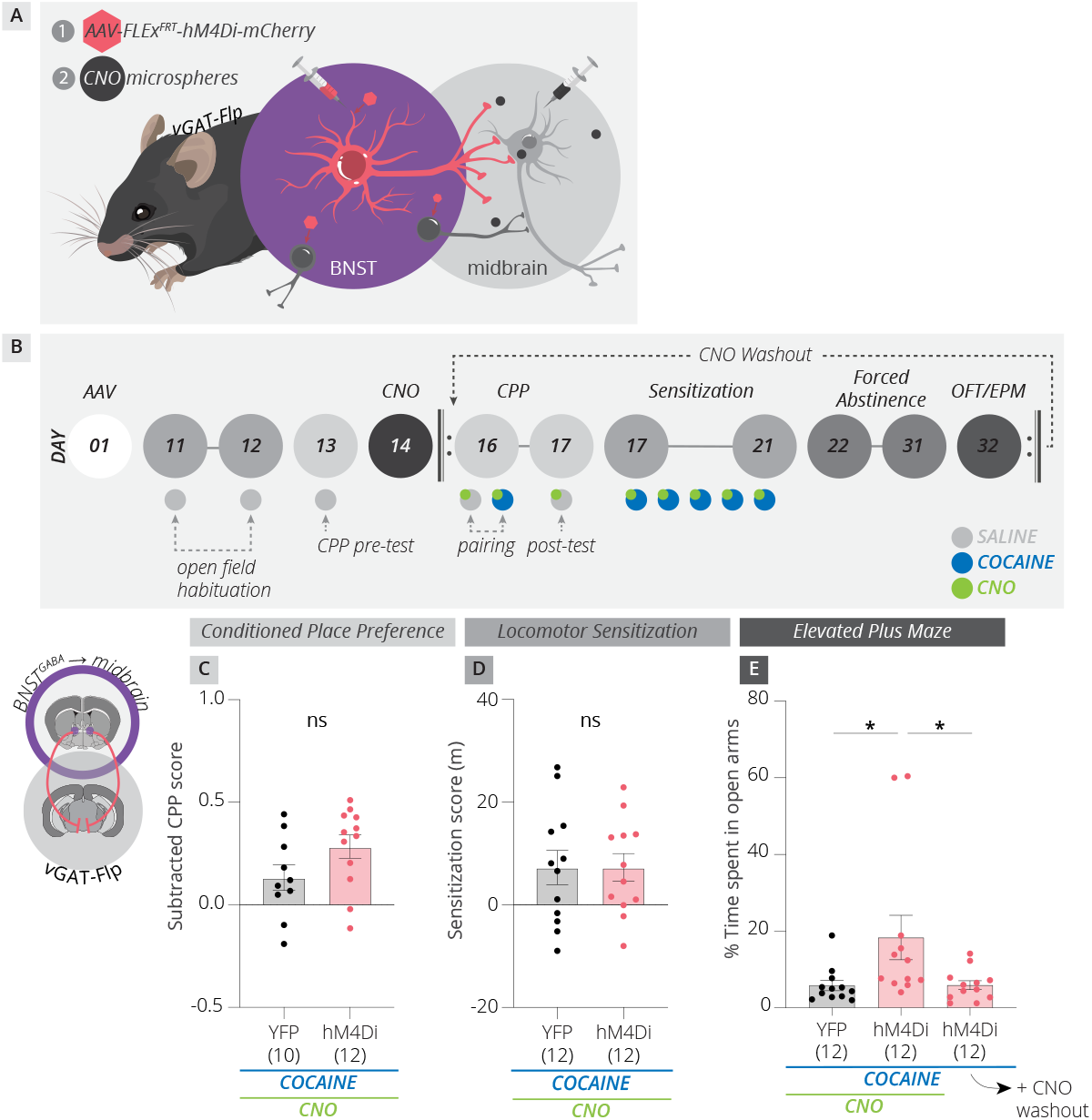
Inhibition of BNST^GABA^ projections to the midbrain prevents development of cocaine withdrawal-induced anxiety. (A) Strategy for terminal inhibition. hM4Di was expressed in BNST^GABA^ cells, and slow-release CNO microspheres were injected into the midbrain to inhibit BNST^GABA^ terminals in the ventral midbrain. (B) Timeline for experiments. The protocol was similar to that performed in Fig. 1 except that all six cocaine injections were given within a 7-day time window when the CNO-releasing beads would be releasing CNO. (C-E) Inhibition of BNST^GABA^ terminals in the midbrain had no effect on CPP (p = 0.09) or sensitization (p = 1.0), but prevented development of anxiety (YFP vs hM4Di, p = 0.023; hM4Di vs washout, p = 0.033; YFP vs washout, p = 0.16).

### Activity in BNST^GABA^→VTA^DA^→Amygdala pathway provides an anxiety signal

Given the importance of the BNST^GABA^→VTA^DA^→Amygdala pathway for development of withdrawal-induced anxiety, we wanted to explore how activity in this pathway related to anxiety behaviors in awake, behaving animals exposed to cocaine. We therefore performed fiber photometry experiments by expressing the calcium-dependent fluorescent protein GCaMP6f in either BNST^GABA^ or VTA^DA^→Amygdala cells, and implanting a chronic fiber over the corresponding cell population. We formed a cocaine administration protocol similar to that performed in Figure 1, although no CNO was administered (Fig. 5A). We first examined the task-linked activity in each cell population during chamber crossings in the CPP pretest and posttest, locomotor initiation and cessation, entries into and exits from the center of the open field, and entries into the closed and open arms of the elevated plus maze (Fig. 5B-C). Both cell populations were minimally active during CPP, locomotion, or the OFT, and were the most active during the EPM task (Fig. 5B-C). Specifically, both BNST^GABA^ and VTA^DA^→Amygdala cells were more active during open arm entries and less active during closed arm entries, with the peak of BNST^GABA^ neuron activity during open and closed arm entries slightly preceding that of VTA^DA^→Amygdala cells (Fig. 5B-C). When either cell population displayed task-related activity (peak z-score greater or less than 0.25; Fig. S5), the activity correlation between BNST^GABA^ and VTA^DA^→Amygdala cells was the highest in the EPM task (r^2^ = 0.85 for the open arm, 0.95 for the closed arm; Fig. 5D, S6). This suggests that activity of these neurons signals anxiety, such as that experienced by entrance into the open arms of the maze. Accordingly, the magnitude of BNST^GABA^ and VTA^DA^→Amygdala cell activation during open arm entries for each mouse correlated with time spent in the open arms for that mouse, with the activity of both populations being inversely related to time spent in the open arms (Fig. 5E-F). Activation or suppression of activity in these cell populations also related to time spent in the center of the open field, but not chamber entries in the CPP task or locomotion (Fig. S7). These results together demonstrate that coordinated activity in these cell populations signals an anxiety state.

**Figure 5:**
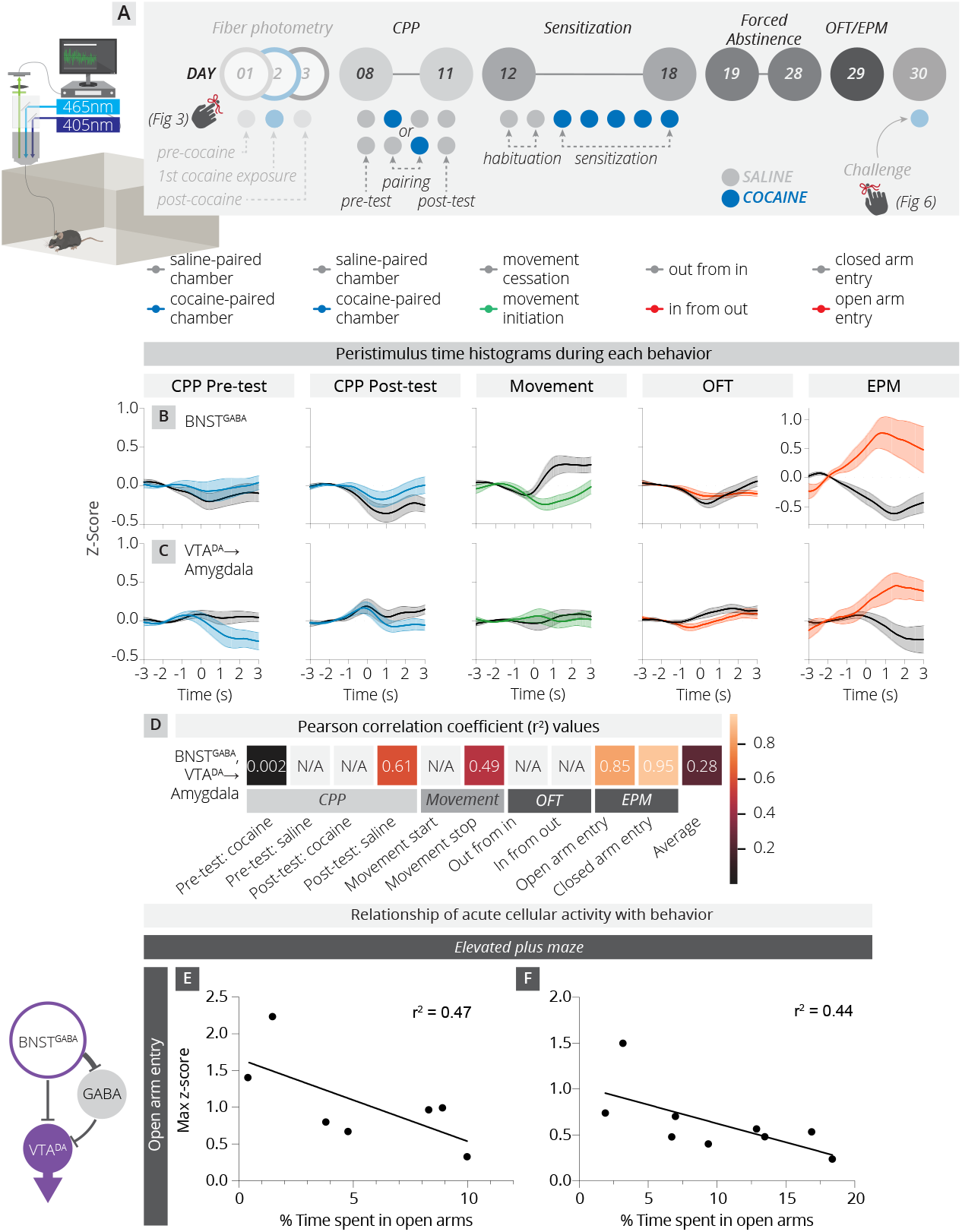
Task-linked cell activity in BNST^GABA^→VTA^DA^→Amygdala pathway is related to anxiety behavior. (A) Schematic and timeline of fiber photometry experiments. The timeline was similar to that used in Figure 1, except that no CNO was given. Portions of the protocol that are the focus of previous (Fig. 3) and subsequent (Fig. 6) figures are denoted. (B-C) Peristimulus time histograms (PSTH) for (B) BNST^GABA^ cells and (C) VTA^DA^→Amygdala cells during five different tasks: chamber crossings into the saline-paired and cocaine-paired chamber in the CPP pre-test and CPP post-test, movement initiation and movement cessation in the open field, entering or exiting the center of the open field in the OFT, and entering the open or closed arm of the EPM. (D) Correlogram showing r^2^ values for activity for BNST^GABA^ cells and VTA^DA^→Amygdala cells in the 0.5 seconds preceding and 1.5 seconds following the identified behavioral event. If neither the maximum or minimum of the z-score of either trace surpassed 0.25 or −0.25, respectively, N/A was applied. (E-F) Correlations of maximum z-scores in (E) BNST^GABA^ and (F) VTA^DA^→Amygdala cells with time in the open arms of the EPM for each mouse.

### Activity in VTA^DA^→Amygdala cells is required for cocaine-primed CPP reinstatement

Withdrawal-related anxiety is one of many factors that contribute to reinstatement of drug taking during abstinence that include re-exposure to drug-paired cues and environmental stressors^35–37^. The amygdala is known to play a central role in cued reinstatement of a variety of behaviors, including drugs of abuse^38–42^. Given the relationship between withdrawal anxiety and relapse, we examined the activity of four different midbrain DA cell population during each cocaine exposure, including a challenge dose of cocaine that was given one day following the EPM and OFT tests after ten days of protracted withdrawal (Fig. 6A). Interestingly, we observed a rhythmic burst of activity following cocaine in VTA^DA^→Amygdala cells that lasted several minutes that was not seen in the other three midbrain DA cell populations examined (Fig. 6B). This activity pattern was present in these cells during all cocaine exposures, but the number of events was about three times greater following the challenge dose than any other cocaine exposure (1.43 events per minute for the first 10 minutes following the challenge dose vs. an average of 0.44 events per minute for all other days; Fig. 6B). This led us to hypothesize that the elevated activity of these cells may be related to reinstatement behavior. To test this possibility, we inhibited VTA^DA^→Amygdala cells during the cocaine priming dose in a CPP reinstatement task (Fig. 6C). *CAV-FLEx^loxP^-Flp* was injected into the amygdala, and Flp-dependent AAVs expressing hM4Di or YFP were injected into the VTA to inhibit VTA^DA^→Amygdala cells or as a control, respectively, as in Fig. 1. Two weeks following virus injection, CPP was established with a single cocaine pairing, as before, and then extinguished over two sessions by allowing the animals to freely explore both chambers of the CPP apparatus in the absence of cocaine. Three days later, CNO was injected in all animals 30 minutes prior to an injection of 7.5 mg/kg cocaine. After cocaine was injected, animals were placed into the CPP apparatus. While YFP-expressing VTA^DA^→Amygdala cells reinstated their CPP, reinstatement was blunted by hM4Di-mediated inhibition of VTA^DA^→Amygdala cells (Fig. 6D). These results demonstrate that activity of VTA^DA^→Amygdala cells during the priming dose is required for cocaine-primed reinstatement of CPP.

**Figure 6:**
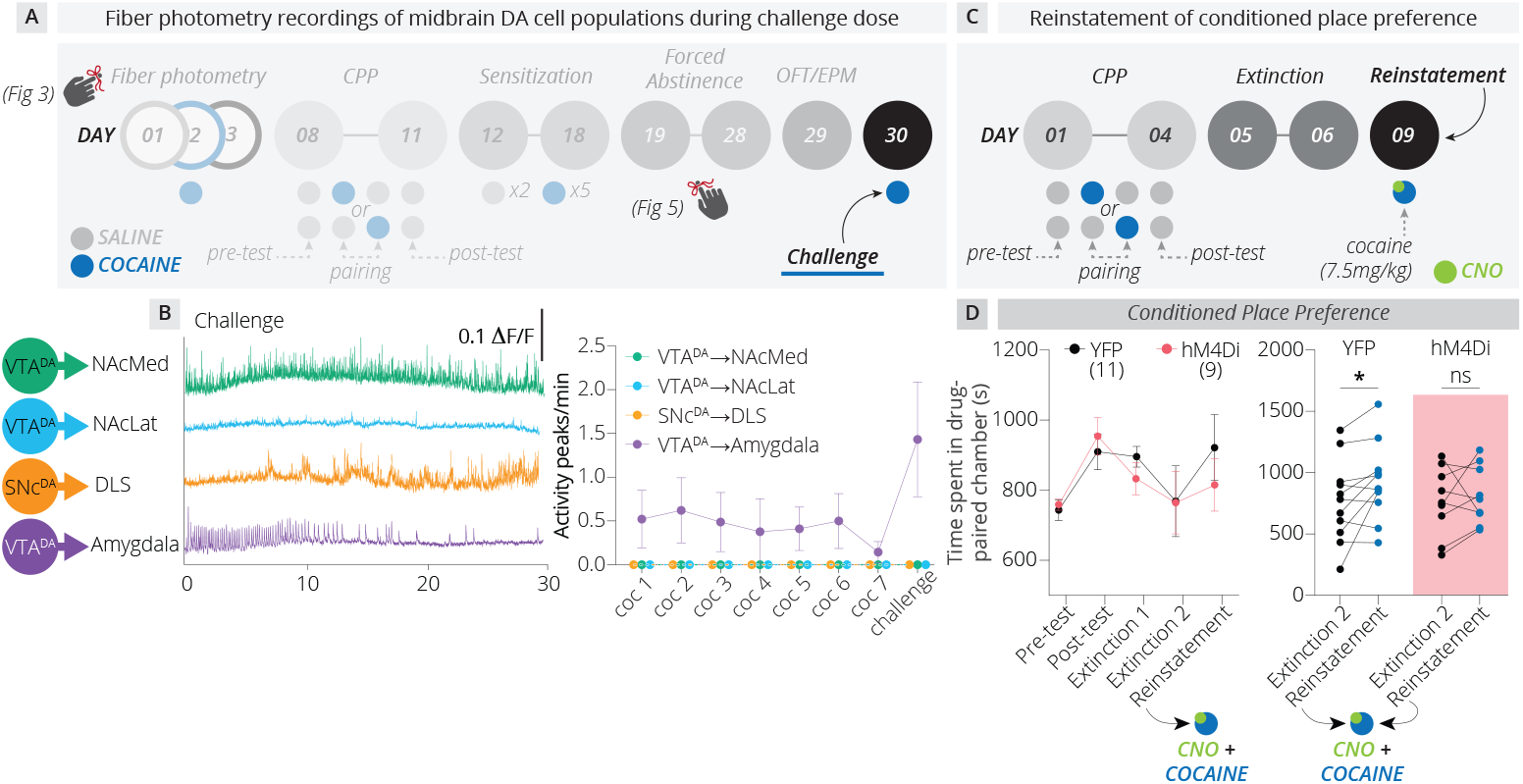
Activity in VTA^DA^→Amygdala cells is required for cocaine-primed CPP reinstatement. **(A)** Timeline for fiber photometry experiments. Mice were the same as those used in Figure 5. **(B)** Representative traces from each of four midbrain DA cell populations in response to a challenge dose of cocaine following ten days of forced abstinence. VTA^DA^→Amygdala cells demonstrated a unique rhythmic activity pattern for approximately the first 10 minutes following cocaine exposure. The frequency of such events for all cell populations during the first 10 minutes following each cocaine dose was quantified. (C) Timeline for CPP reinstatement experiments. CNO was administered only once, 30 minutes preceding the cocaine priming dose. (D) Inhibiting VTA^DA^→Amygdala cells during the priming dose prevented cocaine-induced CPP reinstatement (before/after YFP, p = 0.04; hM4Di, p = 0.57).

### VTA^DA^→Amygdala cell activity is generally required for development of experience-driven anxiety states

Here we have defined the role of the BNST^GABA^→VTA^DA^→Amygdala pathway for the development of cocaine-induced anxiety and reinstatement of CPP. Cocaine is a suitable model experience because it elicits different behavioral adaptations over the course of repeated administration and abstinence, enabling us to dissociate the roles of independent circuit changes in behavioral adaptation. However, the roles these circuits play in behavioral adaptation should be independent of the experience used to elicit them. Therefore, we hypothesized that activity in the BNST^GABA^→VTA^DA^→Amygdala pathway during an experience is required to establish anxiety elicited by that experience. In order to test this hypothesis, we assessed the necessity of activity in VTA^DA^→Amygdala cells for the development of anxiety induced by the model opioid morphine, as well as chronic exposure to a predator odor, a natural stimulus. A combination of *CAV-FLEx^loxP^-Flp* and Flp-dependent AAVs expressing YFP or hM4Di were used to target YFP or hM4Di expression to VTA^DA^→Amygdala cells, as in Figs. 1 and 6. Inhibition of VTA^DA^→Amygdala cells during repeated morphine administration had no effect on CPP or locomotor behavior induced by morphine, but prevented development of anxiety during protracted withdrawal (Fig. 7A-D). We then inhibited these same cells in different animals that were chronically exposed to 2,4,5-trimethylthiazoline (TMT), a constituent of fox urine that is inherently aversive to rodents and elicits anxiety^43^. 4 days of TMT exposure for an hour per day caused long-lasting anxiety in animals injected with YFP relative to animals not exposed to the predator odor, and inhibition of VTA^DA^→Amygdala neurons during odor exposure prevented the development of TMT-induced anxiety (Fig. 7E). These results together demonstrate that activity in the VTA^DA^→Amygdala pathway is generally required for the development of anxiety states and suggest that our method can be generally applied to identify key neuronal circuits that contribute to normal and pathological behavioral adaptations.

**Figure 7:**
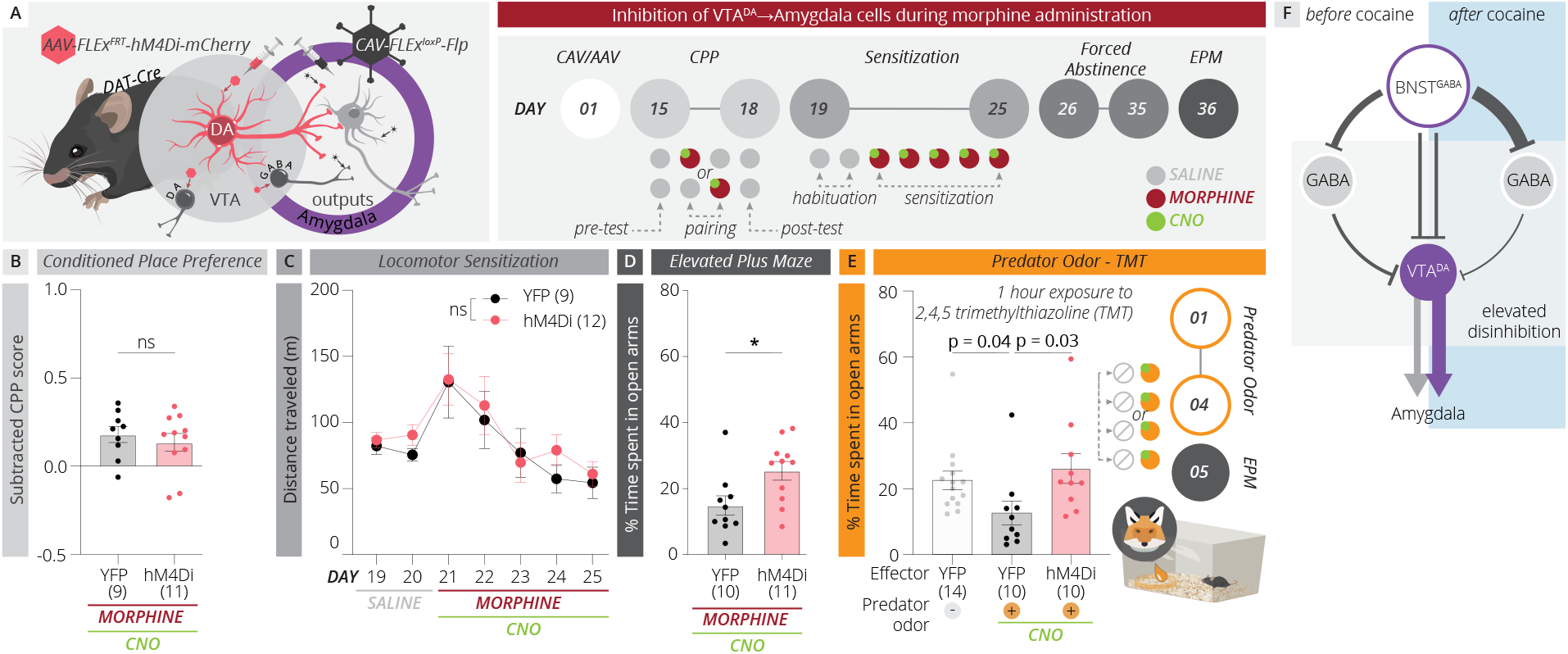
VTA^DA^→Amygdala cell activity is generally required for development of experience-dependent anxiety states. (A) Schematic and timeline of VTA^DA^→Amygdala cell inhibition to test effects on morphine-induced behaviors. (B-D) Inhibition of VTA^DA^→Amygdala cells had no effect on (B) CPP (p = 0.52) or (C) locomotor sensitization (p = 0.34), but it prevented development of (D) anxiety during the forced abstinence period as assessed using the EPM (p = 0.02). (E) Inhibiting VTA^DA^→Amygdala cells prevented anxiety that developed following chronic exposure to TMT (YFP/no PO vs YFP/PO, p = 0.04; YFP/PO vs hM4Di/PO, 0.03; YFP/no PO vs hM4Di/PO, p = 0.49). (F) Proposed circuit model. BNST^GABA^ neurons preferentially inhibit midbrain GABA over midbrain DA cells. Cocaine triggers a long-lasting increase in activity of BNST^GABA^ neurons, which leads to a long-lasting activation of VTA^DA^→Amygdala cells. These changes then contribute to the development and expression of anxiety that occurs following repeated cocaine administration.

## DISCUSSION

In this study, we defined the role of a pathway from BNST^GABA^ cells through midbrain DA neurons to the amygdala in the development of anxiety elicited by both drugs of abuse and natural experiences. Activity in VTA^DA^→Amygdala cells was not required for and had little relationship to cocaine reward or sensitization, but was critical during cocaine exposure for the development of anxiety during protracted withdrawal following repeated cocaine exposure. Using RABV circuit mapping approaches we found that a single exposure to cocaine caused an increase in labeled inputs from the BNST that tracked with an increase in spontaneous activity of these cells 24 hours later without any detectable changes in the number of synapses from BNST^GABA^ cells in the midbrain. Silencing BNST^GABA^ inputs to the midbrain during cocaine administration prevented the development of anxiety, mirroring the effect of downstream VTA^DA^→Amygdala cells. Activity in both BNST^GABA^ cells and downstream VTA^DA^→Amygdala cells was linked to open and closed arm entries in the elevated plus maze, and the extent of activation of each cell type during open arm entries was positively correlated with anxiety. We then showed that activity in VTA^DA^→Amygdala, but not other midbrain DA cells was required for cocaine-primed reinstatement of CPP. Lastly, we showed that activity in VTA^DA^→Amygdala cells was necessary during either morphine administration or predator odor exposure for development of long-lasting anxiety states. The elucidation of how the BNST^GABA^→VTA^DA^→Amygdala pathway selectively orchestrates development of experience-induced anxiety and cocaine reinstatement illuminates a direct link between midbrain and extended amygdala networks in the development of experience-induced anxiety.

The midbrain DA system is critical for the development of drug addiction, and has been shown to play important roles in both the rewarding aspects of drug use as well as of aversive states that develop during withdrawal from drug use. The recruitment of the brain’s stress circuitry, including the extended amygdala, following repeated drug use has been well characterized^44,45^. Withdrawal from repeated drug use leads to a variety of adaptive processes that oppose the acute actions of abused drugs, and these processes lead to the development of an anxiety state during protracted withdrawal that can last for weeks or longer^35,46^. It has been postulated that stress or drug cues induce norepinephrine release into the BNST during protracted drug withdrawal that promotes an anxiety state. This anxiety state in turn may enhance the reward value of drugs via a negative reinforcement mechanism, as temporarily eliminating the anxiety becomes a key driver of drug seeking^35^. It has also been hypothesized that hedonic processing is altered during protracted withdrawal^47–49^, likely through adaptations in midbrain DA cells or their inputs, and that these changes may contribute to relapse/reinstatement. Our results demonstrating the critical role of the BNST and extended amygdala more generally are therefore consistent with previous studies. However, we extend these observations by suggesting that withdrawal-induced anxiety and hedonic changes that lead to reinstatement of drug-seeking behaviors may be orchestrated by a single pathway. We provide an integrated framework by which the BNST and midbrain DA cells work in concert with the amygdala to orchestrate anxiety and reinstatement behaviors induced by withdrawal following cocaine administration and a priming dose of cocaine, respectively. Silencing either BNST^GABA^ inputs to the midbrain, or downstream VTA^DA^→Amygdala cells during cocaine administration is sufficient to prevent the development of anxiety following repeated cocaine administration, demonstrating that activity in these cell populations is critical for long-lasting circuit changes that mediate anxiety. It is likely that cocaine causes long-lasting changes in brain circuits that are normally mediated by these cell populations, and without this activity, the changes that normally mediate anxiety states do not effectively occur. Importantly, our data also highlight that the anxiety that develops following repeated drug exposure is facilitated by circuit elements that are independent of those that mediate drug reward or sensitization, indicating that reward/sensitization and anxiety are driven by different brain circuits.

We also provide further evidence that our RABV mapping platform can be used to identify experience-dependent modifications in neuronal circuits by identifying cell populations that exhibit changes in baseline cellular activity following an experience. Our RABV approach identified a cocaine-induced elevation in BNST^GABA^ cell activity, and inhibition of this projection to the midbrain was sufficient to prevent cocaine-induced anxiety. Interestingly, while a single cocaine exposure is not sufficient to induce long-lasting anxiety, we were still able to detect an elevation in spontaneous BNST cell activity that was important for the development of anxiety later on, suggesting that the RABV technique is sufficiently sensitive to detect early changes in cellular activity that precede behavioral adaptation. In our calcium imaging experiments, we examined cellular activity in all BNST^GABA^ cells, irrespective of output site. Nonetheless were able to detect activity changes in these cells. We used this approach for two reasons: first, we cannot yet perform long-term imaging experiments in BNST^GABA^ cells that selectively provide input to VTA^DA^→Amygdala cells, because RABV is toxic to cells that it infects, and thus likely would alter activity in both VTA^DA^→Amygdala cells and BNST inputs as well as behaviors mediated by these cells. Without RABV-based labeling of connected inputs, we cannot longitudinally map activity in input cells specifically projecting onto spatially intermingled, projection-defined VTA^DA^ cell populations. Defining cells by output (e.g., injection of a retrograde virus into the midbrain) would not maintain connectivity relationships between inputs and defined midbrain DA cell populations. While defining BNST^GABA^ cells by output site may somewhat increase specificity, it would not definitely address this problem. Secondly, we showed that the dominant projection of BNST^GABA^ cells is to the midbrain (Fig. S3), suggesting that more BNST^GABA^ cells projected to the ventral midbrain than any other region. In sum, these results demonstrate that one can first use the RABV method as a screen to identify input brain sites that may contribute to behavioral adaptation. Hits from this screen can then be explored in more detail, for example by using a recombinase line to define the relevant cells in the defined input site. We showed here and in our previous study in the GPe^30^ that this strategy is sufficient to identify cell populations and projections that mediate cocaine-induced behavioral changes.

In addition, while we used cTRIO from VTA^DA^→Amygdala cells to identify cocaine-induced changes in input cell activity, the effect of these input changes on DA cells may not always be direct. It is necessary to consider potential local microcircuit effects, as RABV can identify input changes that may influence multiple cell types in the targeted brain region. In both this and our previous study, the GABAergic cells that we identified using RABV mapping from midbrain DA cells preferentially project onto the GABAergic SNr and thus, elevations of activity from these inputs likely disinhibits midbrain DA cells, leading to an elevation in their activity, as we found here (Fig. 3K) and in our previous study^30^. We believe that this approach is a valuable screening method to enable the dissection of the roles unique circuits play in selective behavioral adaptations, such as the role of the BNST^GABA^→VTA^DA^→Amygdala pathway in DA-dependent establishment of experience-induced anxiety states shown here. Pharmacological intervention for psychostimulant abuse has remained elusive in part because drugs that target the entire DA system often have many off-target effects, including on the brain’s reward system. Elucidation of the selective role of the BNST^GABA^→VTA^DA^→Amygdala pathway in development of anxiety provides specific cellular substrates outside of the canonical DA reward circuits that could be used as targets for the development of addiction therapeutics to reduce negative affect, including anxiety, that develops during withdrawal.

## Acknowledgements

We would like to acknowledge Boris Heifets for providing and assisting with implementation of custom MATLAB scripts for fiber photometry analysis, Yihan Wang for computational assistance, and Greg Corder and Steve Mahler for their critical analysis. This work was supported by the NIH (R00 DA041445, DP2 AG067666), Tobacco Related Disease Research Program (T31KT1437, T31P1426), American Parkinson Disease Association (APDA-5589562), Alzheimer’s Association (AARG-NTF-20-685694), New Vision Research (CCAD2020-002), and the Brain and Behavior Research Foundation (NARSAD 26845) to KTB, T31DT1729 to MH, and T32 GM136624-01 and NSF GRFP to PD.

## METHODS

### Mice and viral procedures

Generation and characterization of the *DAT-Cre*^51^ and *vGAT-Flp* (Jax stock #029591), mouse lines have been described previously. Mice were housed on a 12-h light–dark cycle with food and water *ad libitum.* Males and females from a C57/BL6 background were used for all experiments in approximately equal proportions. All surgeries were done under isoflurane anesthesia. All procedures complied with the animal care standards set forth by the National Institute of Health and were approved by the University of California, Irvine’s Institutional Animal Care and Use Committee (IACUC).

The titers of viruses, based on quantitative PCR analysis, were as follows:

*AAV_5_-CAG-FLEx^FRT^-TC,* 2.6 × 10^12^ genome copies (gc)/ml;
*AAV_8_-CAG-FLEx^FRT^-RABV-G,* 1.3 × 10^12^ gc/ml;
*AAV_DJ_-hSyn-FLEx^FRT^-hM4Di,* 2.9 × 10^13^ gc/ml;
*AAV_DJ_-hSyn-FLEx^FRT^-YFP,* 4.3 × 10^12^ gc/ml;
*AAV_5_-EF1α-fDIO-GCaMP6f,* 7.3 x 10^12^ gc/ml
*AAV_DJ_-hSyn-FLEx^FRT^-mGFP-2A-Synaptophysin-mRuby,* 3.7 × 10^12^ gc/ml;
*CAV-FLEx^loxP^-Flp,* 5.0 × 10^12^ gc/ml;
*RABVΔG,* 5.0 × 10^8^ colony forming units (cfu)/ml

### Drugs

Cocaine was administered at a dose of 15 mg/kg, morphine at 10 mg/kg, and CNO at 5 mg/kg.

### Transsynaptic tracing/cTRIO

cTRIO experiments were performed as previously described^12^, except that a single injection of cocaine (15 mg/kg) or saline was administered one day prior to RABV injection^30^. We injected 500 nL of *CAV-FLEx^loxP^-Flp* unilaterally into amygdala, and during the same surgery, also injected 500 nL of a 1:1 volume mix of *AAV_5_-FLEx^FRT^-TC* and *AAV_8_-FLEx^FRT^-RABV-G* into the VTA. After 13 days, a single injection of cocaine or saline was given IP. A G-deleted, GFP-expressing, EnvA-pseudotyped RABV was injected into the VTA the following day. Animals were sacrificed five days following RABV injection.

In order to identify input changes that occur following an aversive foot shock, in a separate cohort of animals, 13 days following CAV/AAV injection animals were placed into an auditory fear conditioning chamber, as described in the behavior section below. Control animals were administered the tones but no shocks were given. RABV was injected 1 day later, and animals sacrificed five days subsequent, as described above.

Coordinates used for viral injections were (relative to Bregma, midline, or dorsal brain surface and in mm):

NAcMed: AP +1.55, ML 0.7, DV −4.0;
NAcLat: AP +1.45, ML 1.75, DV −4.0;
DLS: AP +0.25, ML 2.5, DV −3.4;
Amygdala: AP −1.43, ML 2.5, DV −4.5;
mPFC: two injections, one at AP +2.15, ML 0.27, DV –2.1 and another at AP +2.15, ML0.27, DV –1.6;
BNST: AP +0.1, ML 0.8, DV −4.0;
VTA: AP −3.2, ML 0.4, DV −4.2;
SNc: AP −3.2, ML 1.2, DV −4.2.

### Histology

#### Fluorescent in situ hybridization (FISH)

FISH was performed as descried by Kishi and colleagues^52^ with modifications. A programmable single-stranded synthesis method called the primer-exchange reaction (PER) was used to amplify the signal.

##### Probe selection

Sequences for 32 *Slc32a1* (vGAT) and 32 *Slc17a6* (vGluT2) probes (39 - 45 bp) covering the whole messenger RNAs were selected from the mm10 reference genome in oligoMiner (https://yin.hms.harvard.edu/oligoMiner/list.html). A 9 bp sequence (tttCATCATCAT) was added onto the 3’ end of all probes as a short DNA primer for PER to generate repetitive PER concatemers. A 42 bp hairpin sequence (ACATCATCATGGGCCTTTTGGCCCATGATGATGTATGATGATG/3InvdT/) was used as a template for PER. As there is no G in the concatemers, the G-C pair in the hairpin template was used as a PER polymerase stopper in the absence of dGTP. A Clean G sequence (CCCCGAAAGTGGCCTCGGGCCTTTTGGCCCGAGGCCACTTTCG) was added to the reaction to deplete trace dGTP contamination. All probes and oligo sequences were ordered from IDT.

##### PER concatemerization

The 32 *Slc32a1* and *Slc17a6* probes were dissolved individually to get 100 μM stock solutions. The probe mixture was prepared by mixing the same volume of all 32 probe solutions. For a typical 100 μl reaction, the following components were included: 10 μl 10x PBS (final concentration for 1x PBS: 137 mM NaCl, 2.7 mM KCl, 10 mM Na_2_HPO_4_, 1.8 mM KH_2_PO_4_, pH 7.4), 10 μl MgSO4 (100 mM), 10 μl dNTPs (10 mM, mixture of dATP, dTTP, dCTP), 10 μl Clean G (2 μM), 10 μl hairpin template (10 μM), 6 μl BST LF polymerase (NEB M0275L), and 36 μl DEPC-treated H_2_O. After the reaction was incubated at 37°C for 15 min to deplete all dGTP contamination, 8 μl probe mixture was added, and then the reaction was kept at 37°C for 3 hrs to generate concatemers with an approximate size of 400-500 bp. The polymerase was inactivated by heating at 80°C for 20 min. A small amount (5 μl) of reaction was loaded into an agarose gel for electrophoresis to check the size of the probe.

##### Brain section preparation

Five days after cTRIO experiments were performed using the amygdala as an output site, mice were transcardially perfused with 1x PBS and then 4% formaldehyde in 1x PBS. Brains were removed and post-fixed in 4% formaldehyde for 24 hrs, and then dehydrated with 30% sucrose. For long-term storage, brains were frozen in an ethanol and dry ice bath and then kept at −80°C. Immediately prior to performing in situ hybridization reactions, brains were cut into 30 μm slices using a cryostat and stored in DEPC treated 1 x PBS. The brain sections covering BNST (AP +0.26 to −0.34) were divided equally into two sets. One set was probed for *Slc32a1* and the other for *Slc17a6.*

##### In situ hybridization

All containers and buffers were either RNase free or treated with DEPC. In 48 wells, brain sections were washed with 1x PBS for 10 min, incubated in 0.1 M triethanolamine (TEA) for 10 min, and then pretreated with 0.25% acetic anhydride in 0.1 M TEA for 10 min. After being washed with 2x SSC for 5 min, brain sections were incubated at 43°C in an oven in wash buffer A (40% formamide, 2x SCC pH 7, 0.2% Tween-20) for 1 hr, and then incubated with prewarmed probe/hyb solution [50 μL PER concatemer in 450 μl hyb solution (40% formamide, 2x SSC pH 7, 0.2% Tween-20, and 10% dextran sulfate)] at 43°C overnight (18-20 hrs). On the second day, brain sections were washed 2×30 min in wash buffer A, 2×45 min in wash buffer B (25% formamide, 2x SCC pH 7, 0.2% Tween-20), and 2×15 min in 2x SSCTw (0.2% tween) at 43°C. Brain sections were then washed 3×5 min in PBSTw (0.2% tween in 1x PBS) at room temperature and then incubated in 1 μM oligo imager (/5Alex647N/tt ATGATGATGT ATGATGATGT /3InvdT/) in 1x PBS solution with 0.2% tween-20 and 10% dextran sulfate at 37°C for 2 hrs. After the incubation, brain sections were washed 4×5 min in PBSTw at room temperature, and then mounted on Superfrost Plus Micro slides (VWR, cat# 48311-703). Once the brain sections were dry on the slide, slides were incubated in PBSTw with DAPI for 30 min, washed 3×5 min with PBSTw, and then Fluoromount-G^TM^ (Invitrogen, cat# 00-4958-02) was applied. Slides were then covered with cover glass (Thermo Scientific, cat# 152460).

##### Imaging and cell counting

Confocal microscopy was performed using an inverted Zeiss LSM700 AxioObserver with a 20x air objective (Plan-Apochromat 20x/0.8). Laser lines used were 488nm and 639nm. Images were z stacks with 20 μm thickness in 2 μm steps. Fiji was used for cell counting. All observed GFP+ neurons in the BNST were quantified as either positive or negative for overlap with *Slc32a1* or *Slc17a6* (639nm) probes.

### List of Probes for FISH

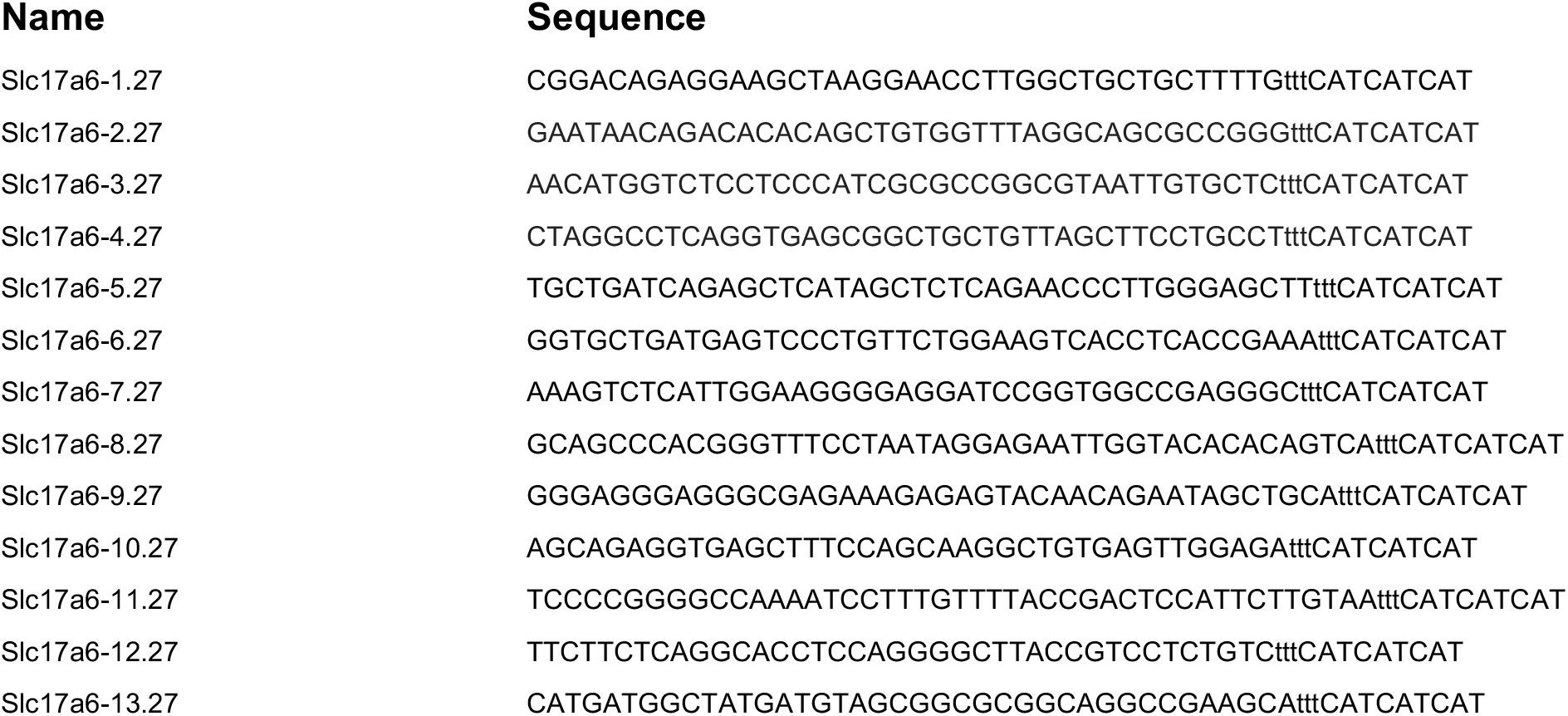
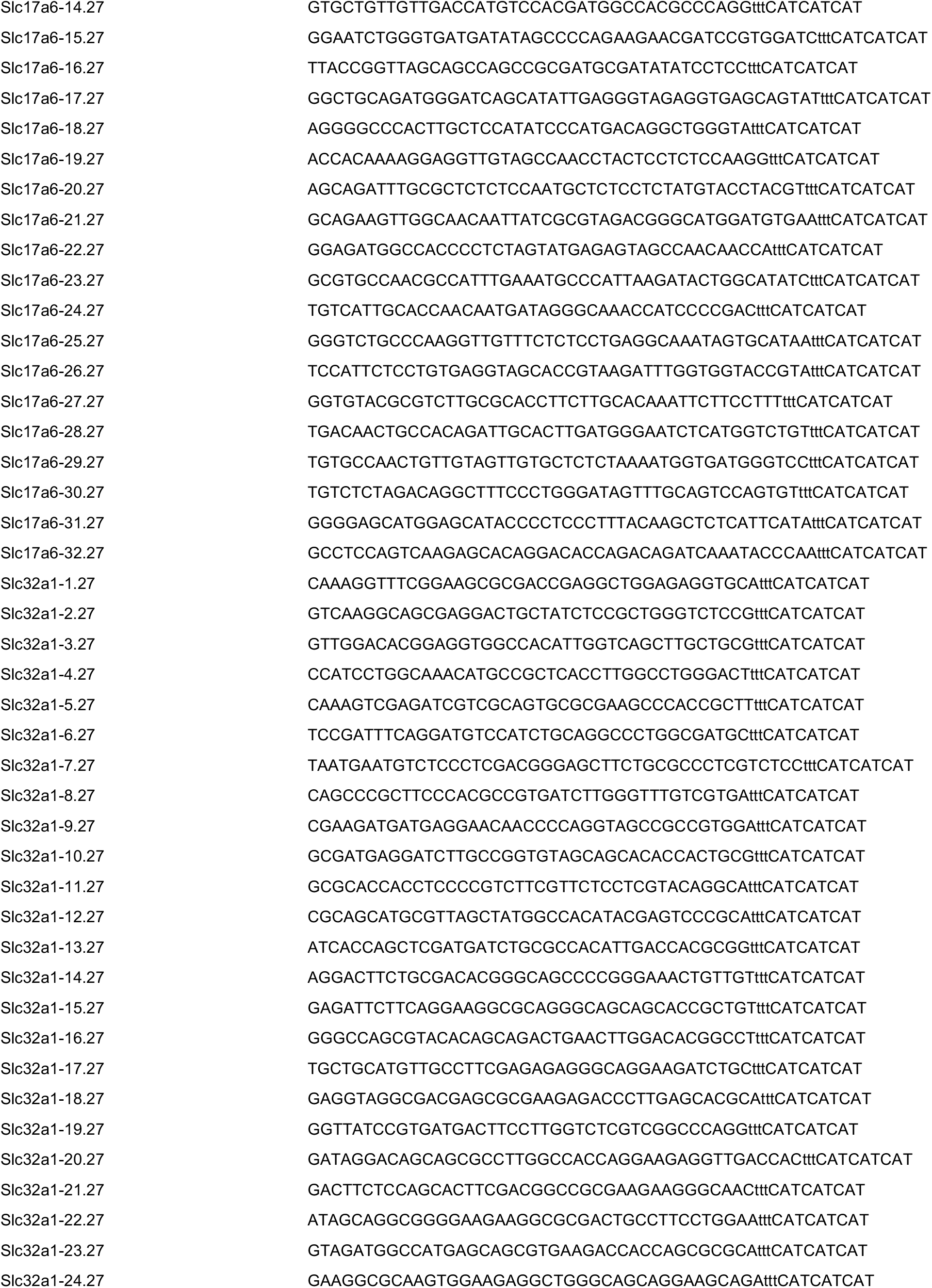
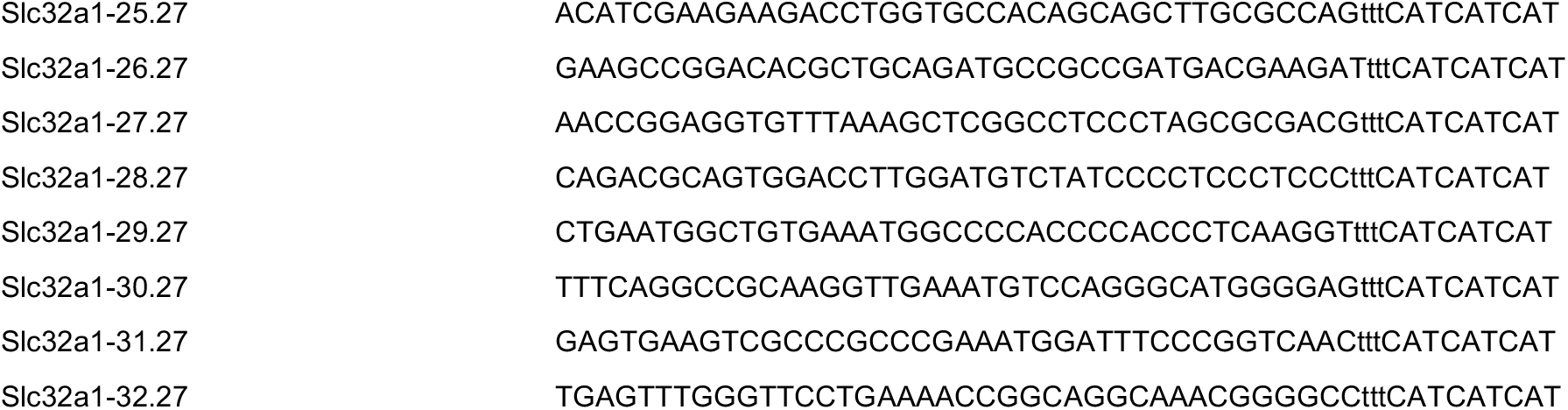

### Immunohistochemistry

The primary antibody chicken anti-GFP (Aves Labs) was used at 1:1,000, rabbit anti-TH (Millipore) at 1:1,000, and rat anti-mCherry (ThermoFisher) at 1:2,000. All secondary antibodies (Donkey anti-chicken AlexaFluor488, donkey anti-rat AlexaFluor 555, donkey anti-rabbit AlexaFluor 647) were used at a concentration of 1:250.

### Axonal arborization mapping from BNST^GABA^ cells

Axonal tracing experiments were performed as previously described^12^. We injected 500 nL of *AAV_DJ_-hSyn-FLEx^FRT^-mGFP-2A-Synaptophysin-mRuby* unilaterally into the BNST. After 2 months, animals were sacrificed, brains were cut with a thickness of 60 μm and immunolabeled for GFP to enhance signal. Five brains were quantified.

Quantifications of puncta from BNST^GABA^ neurons in the midbrain was performed as previously described^30^. Briefly, animals were injected with saline or cocaine 24 hours prior to being sacrificed. Floating sections (60 μm) were stained using anti-mCherry and anti-GFP antibodies. Sections were imaged on a Zeiss 700 confocal microscope using a 63x objective, with image stacks containing 30 sections at 0.44 μm intervals using 2x averaging. Three images were taken of puncta in the SNr and PVT of each brain. Images were analyzed using Imaris (Bitplane). The surface function was used to obtain the volume of mGFP+ neurites and mRuby+ puncta, while the spots function was used to estimate the number of mRuby+ puncta. Data from the three slices from the SNr and PVT were averaged for each brain. Measurements from the SNr were normalized to those from the PVT.

### Fiber photometry

Fiber photometry experiments were performed as previously described^30^. To measure activity in BNST^GABA^ cells, 500 nL of *AAV_5_-fDIO-EF1_α_-GCaMP6f* was injected into the BNST. A 400 μm diameter, 0.48NA optical fiber (Doric Lenses) was implanted at the same location. For VTA^DA^→Amygdala neurons, *AAV_5_-fDIO-EF1_α_-GCaMP6f* was injected into the VTA, 500 nL of *CAV-FLEx^loxP^-Flp* was injected into the amygdala, and fibers were implanted over the VTA. Implants were placed approximately 0.3 mm above the site of viral injection and were secured to the skull with metal screws (Antrin Miniature Specialists), Metabond (Parkell), and Geristore dental epoxy (DenMat). Mice were allowed to recover for at least 4 weeks before experiments.

Fiber photometry recordings were made using previously described equipment^14^. Mice were run through an injection/behavior timeline shown in Fig. 4a. In brief, 405 nm and 465 nm excitation light were controlled via a RZ5P real-time processor (Tucker Davis Technologies) using Synapse software, and were used to stimulate Ca^2+^-dependent and isosbestic emission, respectively. All optical signals were band pass filtered with a Fluorescence Mini Cube FMC4 (Doric) and were measured with a femtowatt photoreceiver (2151; Newport). Signal processing was performed with MATLAB (MathWorks Inc.). Signals were first motion corrected by subtracting the least squares best fit of the control trace to the calcium signal. Data points containing large motion artefacts were then manually removed. To assess neural activity, we quantified the time spent above threshold, which was set at 2.91 times the median absolute deviation (MAD) of each day’s recording, a value that equates to the 95% confidence interval for Gaussian data^33^.

### Peri-stimulus time histograms (PSTHs)

PSTHs were generated using timestamps corresponding to certain behavioral events. Mouse behavior was recorded using Biobserve, which returns a frame-by-frame record of coordinates corresponding to the nose, center of body, and the base of the mouse’s tail. After defining the borders of each behavior arena within Biobserve, a custom Python script was used to generate a list of timestamps corresponding to selected behavioral events. These events are as follows: for both the CPP pre-test and post-test, the moment the mouse enters either the saline-paired or cocaine-paired chamber; for locomotion, the moment the mouse initiated or ceased movement; for OFT, the moment the mouse either entered or left the center zone (where the center zone is defined by a square 1/3 the area of the total arena); for EPM, the moment the mouse entered either the open or closed arms of the maze. The coordinate corresponding to the mouse’s body center was used for all calculations. For locomotion, motion was calculated by taking the distance the center body coordinate moved from frame to frame. Because of some inherent variability in Biobserve’s estimation of the center coordinate (even when the mouse was not moving), velocity below 1.25cm/s was used as the threshold for no movement. A second MATLAB script aligned each behavioral timestamp with the raw photometry trace and resampled the Biobserve coordinates to match the sampling rate of the fiber photometry data. The PSTH curve was then generated by charting the mean Z-score during the three-second pre- and post-event time intervals centered around each behavioral event.

For constructing correlograms, Pearson’s correlation was used to measure synchronization of PSTH curves per region pair for each test and measurement during the time interval of 0.5 seconds preceding and 1.5 seconds following each event. Comparisons where neither PSTH reached a magnitude of 0.25 were dropped (N/A) to focus on synchronous cellular activity. r^2^ and p-values were visualized in heatmaps.

### Behavioral assays

#### CPP

To test for drug-induced CPP, animals were first tested in a single drug pairing, two-chamber CPP test. Each chamber was given different wall contexts. On the first day, animals were initially placed into the right chamber, and allowed to freely explore both chambers for thirty min (pre-test). On the second day, animals were saline-conditioned to the left side, and on the following day, cocaine (or morphine)-conditioned to the right side. Drug conditioning was counterbalanced across the mice. On the fourth day, animals were again initially placed into the right chamber, and allowed to explore freely (post-test). For animals injected with CNO, 5 mg/kg CNO was injected thirty min before the beginning of both the cocaine and saline pairings. CPP scores were computed as the subtracted CPP score, which equals time spent in the drug paired chamber [(posttest-pretest)/posttest]. Each session was 30 min.

#### Sensitization

The following week, to test sensitization, animals were habituated to open field boxes equipped with motion tracking for two days (receiving saline injections before each session). Boxes contained polka dotted contexts on the walls for the duration of the sensitization testing. Animals were then injected with cocaine (or morphine) immediately before entry into the open-field boxes, for five consecutive days, for 30 min each.

#### OFT/EPM

After waiting ten days following the final drug injection, we then tested the animals in the open field test and elevated plus maze for anxiety behaviors. For the OFT the time spent in the center square (1/3 of the total area of the arena) during a five-minute test period was calculated. The following day, animals were tested in the EPM. Animals were placed at the end of one of the open arms, facing outwards, at the beginning of the trial. The percentage of time spent in the two open arms during the five-minute testing period was quantified. OFT experiments were performed at an illuminance level ~1000 lux, and EPM ~25 lux.

For fiber photometry experiments, behavioral assays were performed exactly as described above. For chemogenetic inhibition experiments to test the necessity of defined cell populations in behavioral adaptation, to inhibit projection-defined subsets of midbrain DA cells (Fig. 1, Fig. S1), we used a viral-genetic intersectional method. We injected 500 nL of *CAV-FLEx^loxP^-Flp* bilaterally into the NAcMed, NAcLat, DLS, or amygdala, or 2 μL into the mPFC to target DA neurons projecting to each of these sites. During the same surgery, 500 nL of *AAV_DJ_-hSyn-FLEx^FRT^-hM4Di* (or YFP) was injected bilaterally into the ventral midbrain. Animals were allowed 2 weeks to recover. 5 mg/kg CNO was injected IP 30 min before each injection of 15 mg/kg cocaine (or saline during CPP pairing and day 3 of locomotor habituation to test for CNO effects on basal locomotor activity).

For inhibition of inputs to the midbrain (Fig. 4), we injected 500 nL of *AAV_DJ_-hSyn-FLEx^FRT^-hM4Di* into the BNST of *vGAT-Flp* mice. To inhibit activity in these cells, we used CNO microspheres, as reported previously^30^. CNO microspheres were synthesized to enable slow release of CNO after a single infusion. Degradex PLGA CNO microspheres were custom ordered from Phosphorex. Beads of target mean diameter 1 μm were dissolved in 0.5% trehalose at a concentration of 5 mg micro-sphere per ml. The estimated CNO loading efficiency was 5%, the estimated burst release was 50%, and the estimated release time was 7 days. The target concentration of CNO release at a steady state was 100 pg/h^34,53^.

Given the estimated release time of 7 days, we performed experiments under a condensed timeline to test CPP and sensitization while giving all cocaine doses within a 7-day period. Briefly, animals were habituated to the open field boxes for 2 days before starting CPP experiments. Animals then underwent their CPP pretests. On the following day, 500 nL of CNO microspheres were bilaterally injected into the midbrain. After allowing an additional day for recovery, animals were saline paired with one chamber, followed by a cocaine pairing with the opposite chamber on the same day. The following day, animals were first tested for CPP in the CPP boxes, then were administered cocaine in the open field boxes to start sensitization on the same day. Animals then received 1 dose of cocaine and their locomotion was quantified in the open field for the following four days. Animals then underwent ten days of forced abstinence followed by testing in the OFT and EPM. If significant effects were observed through chemogenetic perturbation of inputs from the targeted cells, we then performed CPP experiments starting with a pretest on day 1, saline and cocaine pairing on day 2, followed by a posttest on day 3, allowing us to perform the CPP over 3 days, as before. This was followed by the first day of sensitization experiments in the open field chamber the same day as the CPP posttest was conducted in order to mirror the first round of experimentation. The rest of the protocol was repeated verbatim.

Morphine CPP, sensitization, and EPM/OFT behavioral assays (Fig. 7A-D) were conducted exactly as for cocaine. 10 mg/kg morphine doses were administered each time, preceded 30 min by 5 mg/kg CNO according to the same protocol as for cocaine.

##### Reinstatement

For cocaine CPP reinstatement experiments in Fig. 6C-D cocaine CPP following a single drug pairing was performed as described above. Following CPP testing, CPP was then extinguished by placing the animals in the same CPP chambers for 2 subsequent days with no drug pairings. After a further 2-day break, animals were injected with CNO, followed 30 min later with a half normal dose of cocaine (7.5 mg/kg), and placed into the CPP chamber, where the time spent in the drug-paired chamber was assessed. Rather than calculating a subtracted CPP score, we report the total time spent in the drug-paired chamber during each 30-minute session.

#### Predator odor stress

2,4,5 trimethylthiazoline (TMT; Sigma), a constituent of fox urine that is innately aversive to rodents, was used for the predator odor. Scent was prepared by mixing 200 μL of TMT into 150g standard mouse bedding and then dividing the scented bedding into 5 10-cm plastic petri dishes sealed with lab tape and perforated to allow the scent to escape. This preparation exposes each animal to approximately 40 μL TMT. Each experimental animal (YFP and hM4Di) was exposed to TMT in a standard mouse cage (36cm L x 20cm W x 13cm D) for 1 hr per day at the same time each day, for 4 consecutive days. Separate control (YFP) mice were handled 2 min per day by the experimenter for each of the 4 days and placed into new cages for 1 hr but were not exposed to the predator odor. On the day following the final odor exposure, experimental and control mice were removed from the vivarium at the same time as the odor exposure was performed in previous days and moved to a different room for the EPM test.

#### Auditory fear conditioning

Mice were first habituated to the auditory fear conditioning chambers. Mice were individually placed in the chamber (Coulbourn Instruments) located in the center of a sound attenuating cubicle. The conditioning chamber was cleaned with 10% ethanol to provide a background odor. A ventilation fan provided a background noise at ~55 dB. After a 2 min exploration period, three 2 kHz, 85 dB tones were played, for 30s each, with a 90s interval between them. Following these initial tones, three subsequent tones were pained with a 1s, 0.75 mA foot shock. The foot shocks co-terminated with the tone. The mice remained in the chamber for another 60s before being returned to the home cages. RABV injections were then performed the following day.

#### Data analyses and statistics

All statistics were calculated using GraphPad Prism 9 software. Statistical significance between direct comparisons was assessed by unpaired or paired t-tests. When multiple conditions were compared, one- or two-way ANOVAs were first performed, as appropriate, and if significant differences were identified, t-tests were then performed for each individual comparison. Multiple comparisons corrections were performed when multiple such t-tests were being performed, and significance was assessed using the Holm-Bonferroni method. In conditions where multiple comparisons were performed and the results were still considered significant, asterisks were presented corresponding to the original p-values. Where the differences were not significant when considering multiple comparisons (e.g., Fig. 1I), the raw, uncorrected p-values are provided on the graph. Dot plots presented throughout the manuscript include a bar representing the mean value for each group. Error bars represent s.e.m. throughout. For all figures, ns P > 0.05, * P ≤ 0.05, ** P ≤ 0.01, *** P ≤ 0.001, **** P ≤ 0.0001.

---

**Figure S1:**
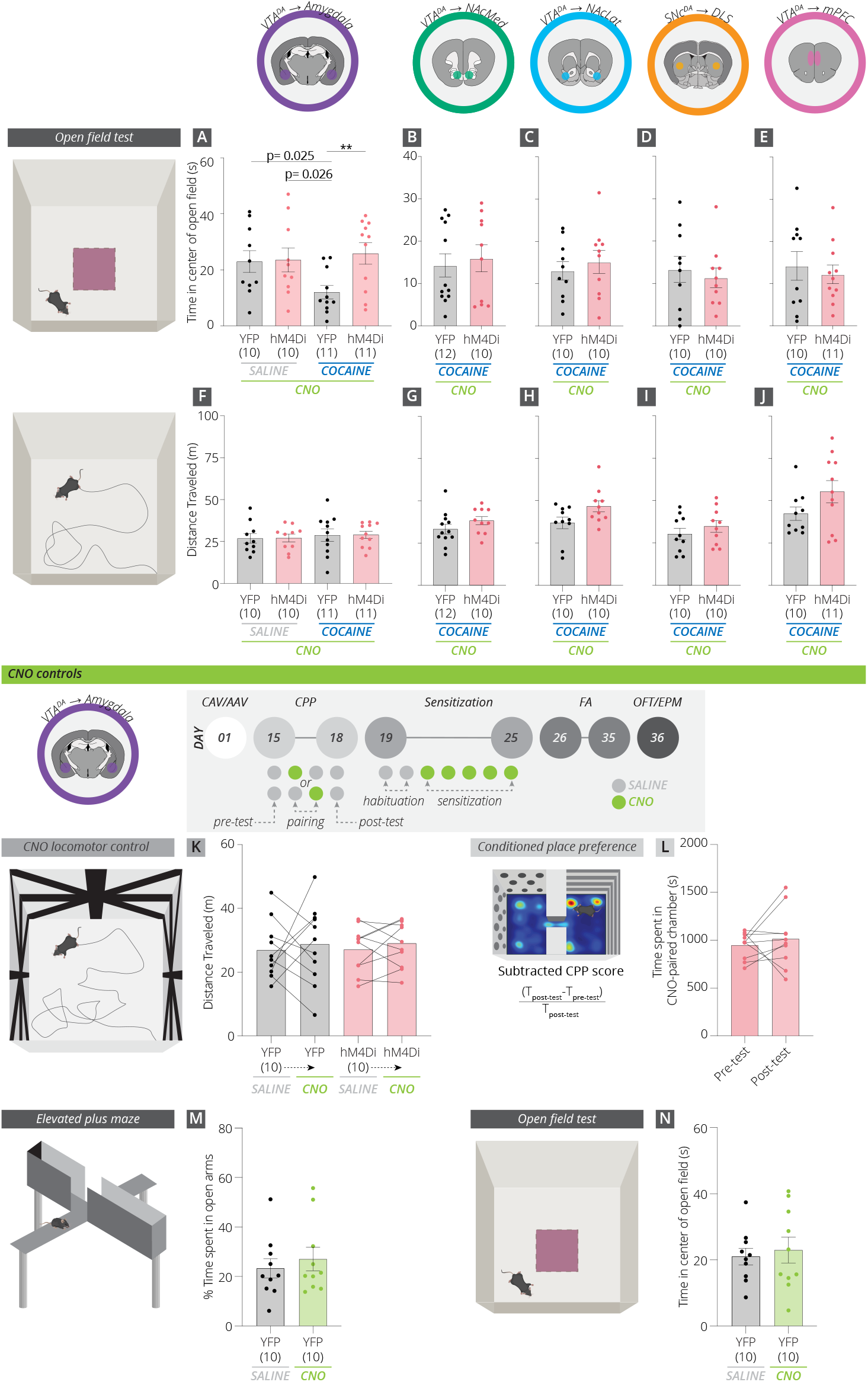
Effects of inhibition of midbrain DA cell populations during cocaine administration on time spent in the center of the open field, plus locomotion and CNO controls. (A) Animals tested during the protracted withdrawal period following repeated cocaine injections spent less time in the center of the open field than saline-treated controls, an effect blocked by inhibition of VTA^DA^→Amygdala cells during cocaine administration (uncorrected p-values, YFP saline vs YFP cocaine, p = 0.0032; YFP saline vs. hM4Di saline, p = 0.53; YFP saline vs hM4Di cocaine, p = 0.79; YFP cocaine vs. hM4Di saline, p = 0.0064; YFP cocaine vs. hM4Di cocaine, p = 0.0098; hM4Di saline vs. hM4Di cocaine, p = 0.73). (B-E) hM4Di-mediated inhibition had no effect on the time spent in the center of the open field for VTA^DA^→NAcMed (p = 0.70), VTA^DA^→NAcLat (p = 0.55), SNc^DA^→DLS (p = 0.62), or VTA^DA^→mPFC cells (p = 0.62). (F-J) Overall locomotion during the OFT for each condition. No significant differences were observed (VTA^DA^→ Amygdala, p = 0.18, VTA^DA^→NAcMed, p = 0.21; VTA^DA^→NAcLat, p = 0.10, SNcDA→DLS, p = 0.36, VTA^DA^→mPFC, p = 0.11). (K) No effects on basal locomotor activity were observed upon CNO administration in animals expressing YFP or hM4Di in VTA^DA^→Amygdala cells (saline administered on Day 20, saline + CNO on Day 21, Fig. 1A: YFP, p = 0.85; hM4Di, p = 0.56). (L) Pairing one chamber of the CPP apparatus with CNO instead of cocaine in hM4Di-expressing animals did not result in a place preference or aversion (Days 15-18, above timeline; p = 0.52). (M-N) Animals treated with 6 doses of CNO and no cocaine showed no long-lasting anxiety effects as YFP-expressing, CNO-treated animals had similar levels of anxiety as control YFP-expressing, saline-treated animals in the EPM (p = 0.54) and OFT (p = 0.68). YFP-expressing, CNO-treated animals are the same as those shown in Fig. 1D and Fig. S1A.

**Figure S2:**
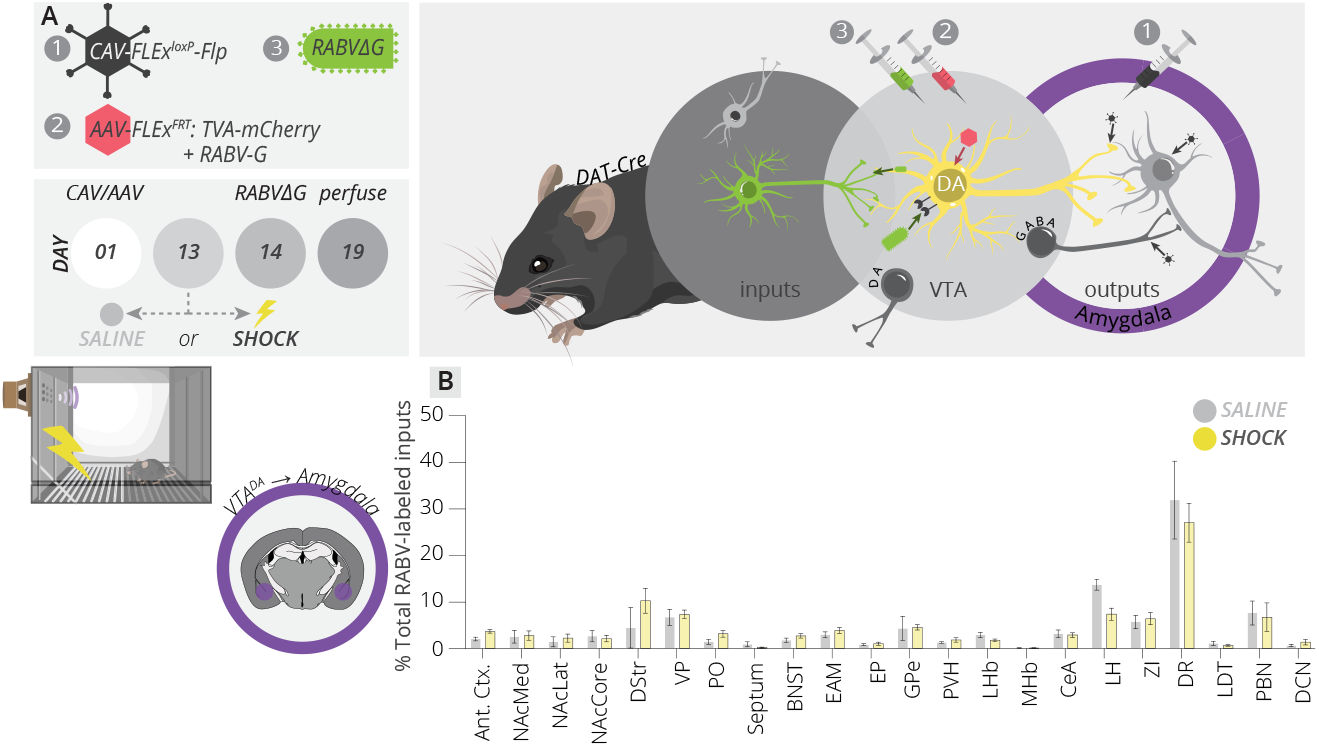
RABV mapping from VTA^DA^→Amygdala cells after receiving three foot shocks. (A) Schematic of cTRIO protocol from VTA^DA^→Amygdala cells with three foot shocks preceding RABV injection by one day. (B) No changes in input labeling were observed onto VTA^DA^→Amygdala cells.

**Figure S3:**
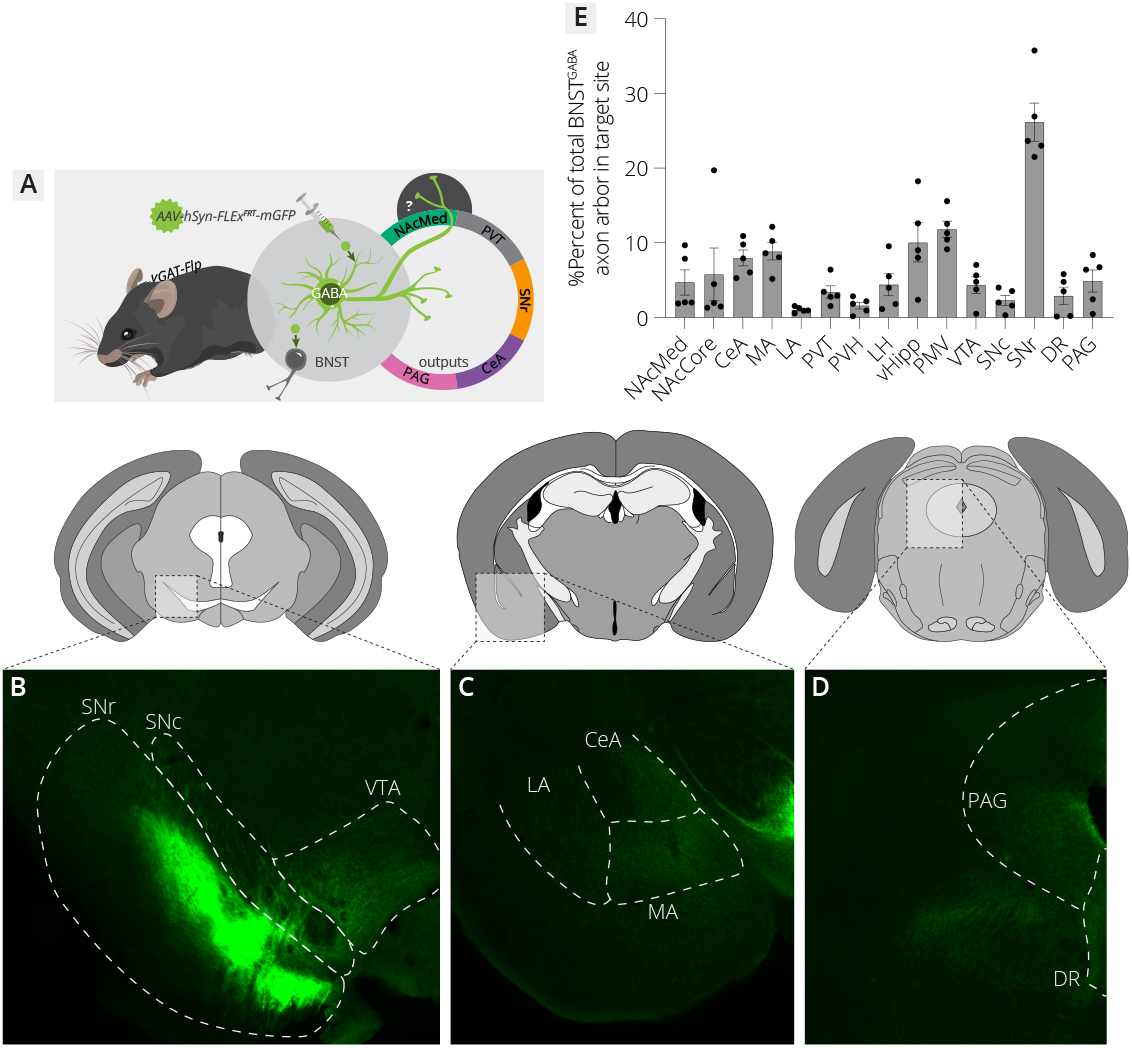
Mapping global outputs of BNST^GABA^ cells. (A) Viral mapping strategy. mGFP was expressed in all BNST^GABA^ cells, and the expression of mGFP in output sites throughout the brain was quantified. (B-D) Sample images of axon labeling in the (B) ventral midbrain, (C) amygdala, (D) periaqueductal gray. (E) Quantification of the percentage of total axons quantified in the target brain region relative to total axons quantified. The SNr received the strongest projection from BNST^GABA^ cells. Region abbreviations: CeA, central amygdala; MA, medial amygdala; LA, lateral amygdala; PVT, paraventricular nucleus; PVH, paraventricular hypothalamus; LH, lateral hypothalamus; vHipp, ventral hippocampus; PMV, ventral premammillary nucleus; DR, dorsal raphe; PAG, periaqueductal gray.

**Figure S4:**
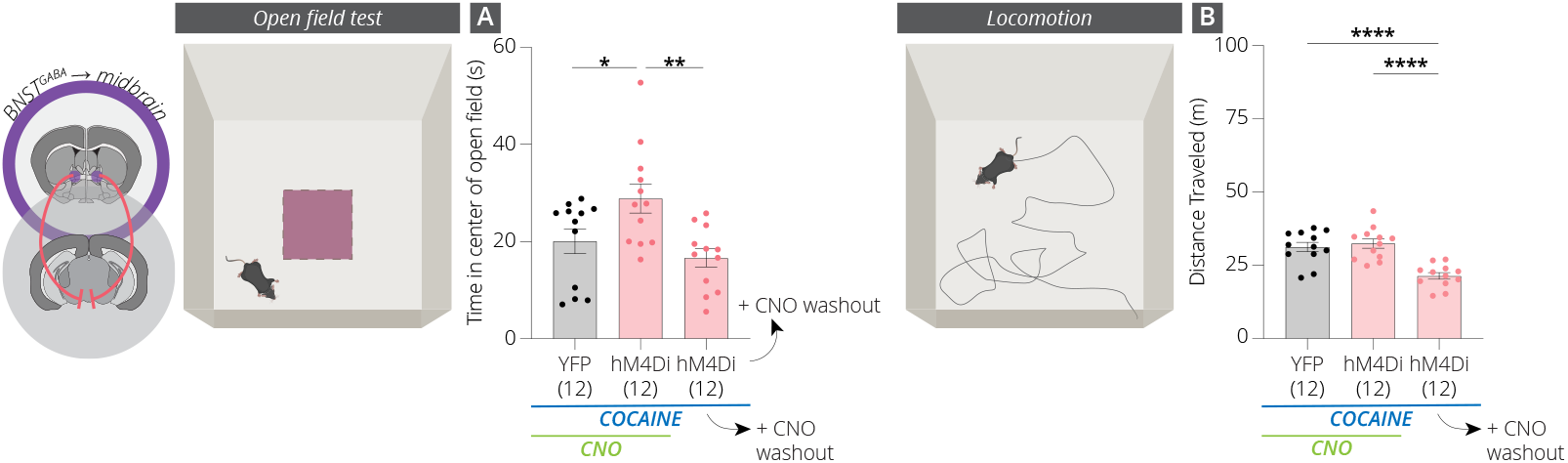
Effects of BNST^GABA^ terminal inhibition in the midbrain during cocaine administration on time spent in the center of the open field and locomotion in the open field. (A) Time spent in the center of the open field (YFP vs hM4Di, p = 0.04: hM4Di vs washout, p = 0.002: YFP vs washout, p = 0.29). (B) Overall locomotion during the OFT (YFP vs hM4Di, p = 0.61: hM4Di vs washout, p < 0.0001: YFP vs washout, p < 0.0001).

**Figure S5:**
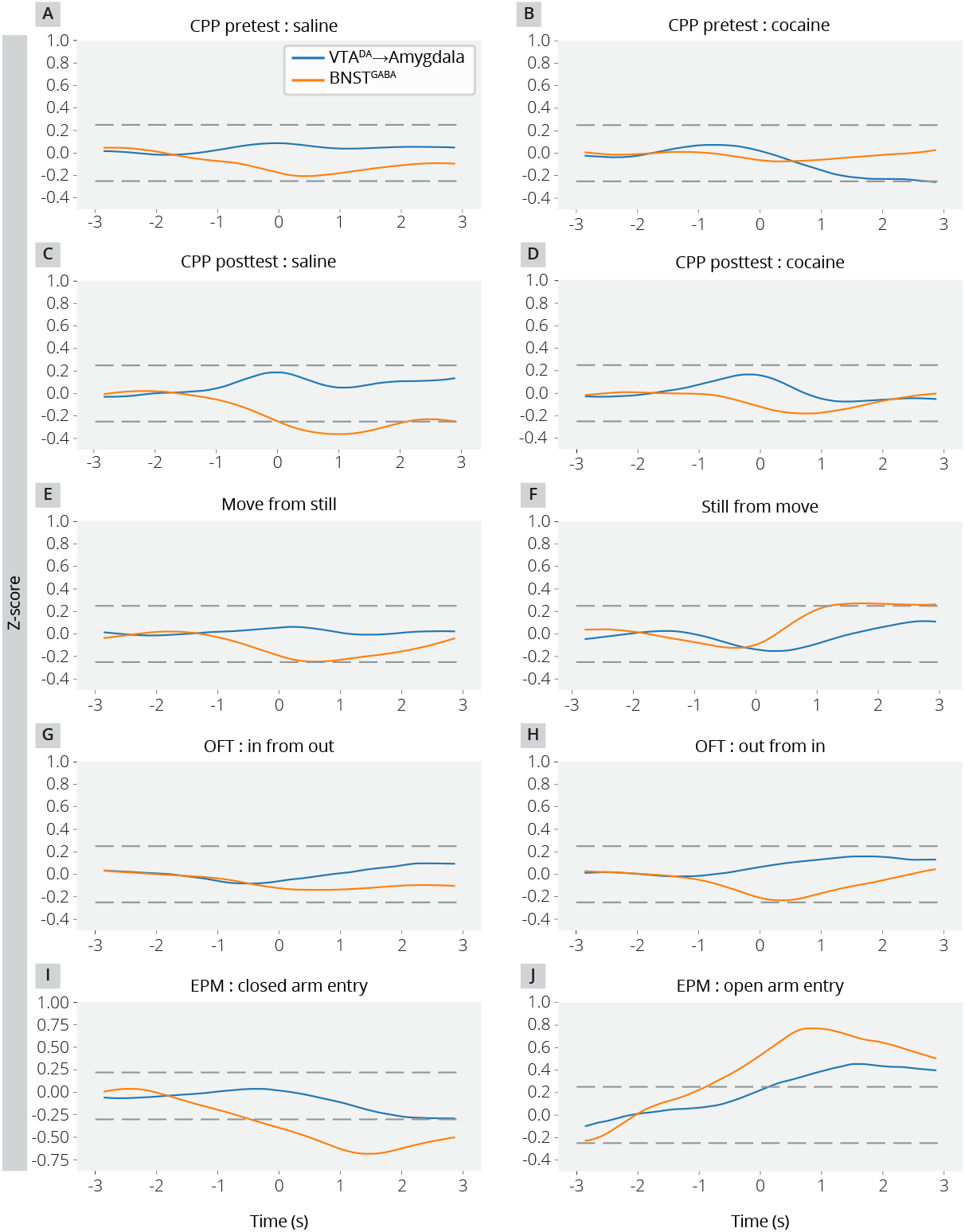
Paired BNST^GABA^ and VTA^DA^→Amygdala cell traces for each behavior assessed. Results were used to generate r^2^ values (Fig. 5D) and p-values (Fig. S6) for correlograms.

**Figure S6:**
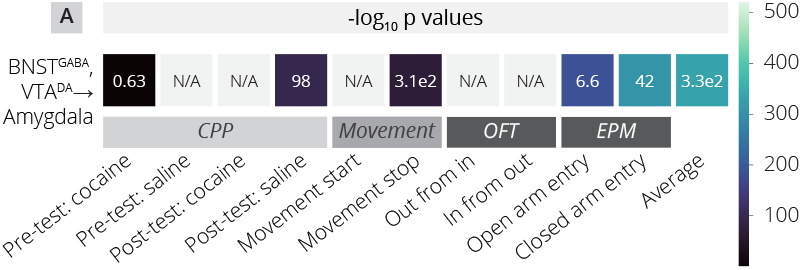
Correlogram showing p-values for paired activity between BNST^GABA^ and VTA^DA^→Amygdala cells. These p-values correspond to r^2^ values shown in Fig. 5D.

**Figure S7:**
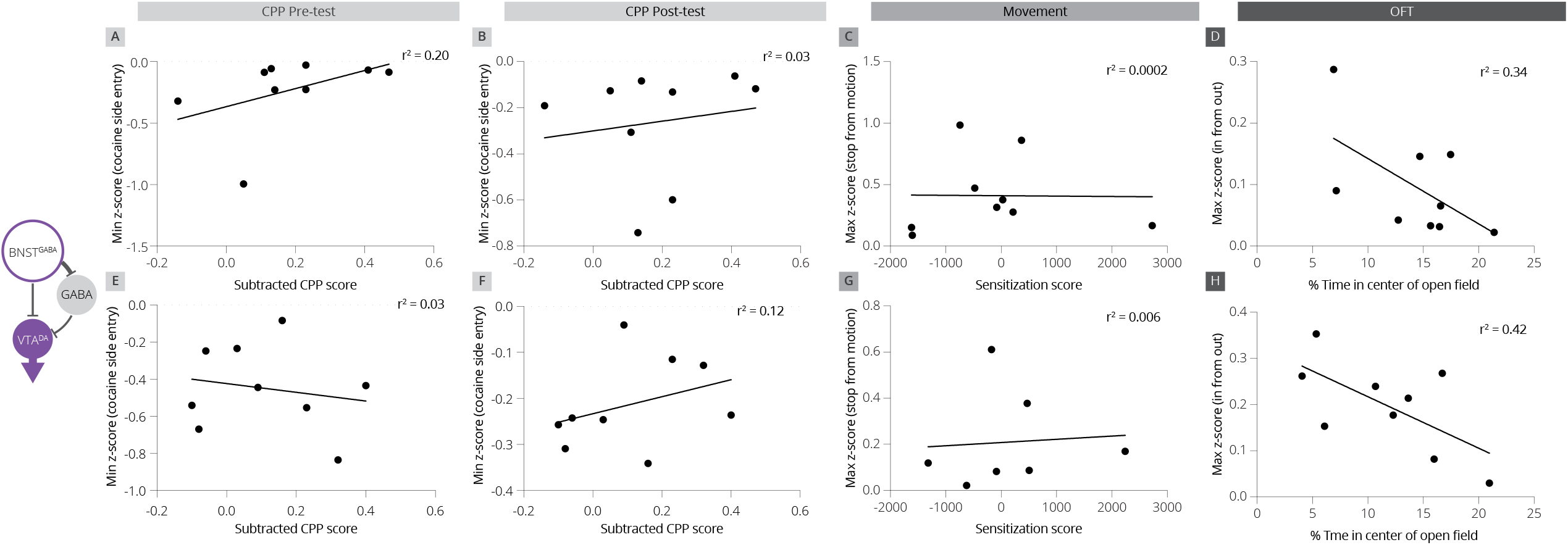
Correlation plots of cellular activity in BNST^GABA^ and VTA^DA^→Amygdala cells with various behaviors. (A) Correlation of the minimum z-score in BNST^GABA^ cells following cocaine-paired side entry with the subtracted CPP score during the CPP pre-test. (B) Correlation of the minimum z-score in BNST^GABA^ cells following cocaine-paired side entry with the subtracted CPP score during the CPP post-test. (C) Correlation of the maximum z-score in BNST^GABA^ cells following locomotion cessation. (D) Correlation of the maximum z-score in BNST^GABA^ cells when entering the center of the open field. (E) Correlation of the minimum z-score in VTA^DA^→Amygdala cells following cocaine-paired side entry with the subtracted CPP score during the CPP pre-test. (F) Correlation of the minimum z-score in VTA^DA^→Amygdala cells following cocaine-paired side entry with the subtracted CPP score during the CPP post-test. (G) Correlation of the maximum z-score in VTA^DA^→Amygdala cells following locomotion cessation. (H) Correlation of the maximum z-score in VTA^DA^→Amygdala cells when entering the center of the open field.

